# Optineurin-facilitated axonal mitochondria delivery promotes neuroprotection and axon regeneration

**DOI:** 10.1101/2024.04.02.587832

**Authors:** Dong Liu, Hannah C. Webber, Fuyun Bian, Yangfan Xu, Manjari Prakash, Xue Feng, Ming Yang, Hang Yang, In-Jee You, Liang Li, Liping Liu, Pingting Liu, Haoliang Huang, Chien-Yi Chang, Liang Liu, Sahil H Shah, Anna La Torre, Derek S. Welsbie, Yang Sun, Xin Duan, Jeffrey Louis Goldberg, Marcus Braun, Zdenek Lansky, Yang Hu

## Abstract

Optineurin (OPTN) mutations are linked to amyotrophic lateral sclerosis (ALS) and normal tension glaucoma (NTG), but a relevant animal model is lacking, and the molecular mechanisms underlying neurodegeneration are unknown. We found that OPTN C-terminus truncation (OPTNΔC) causes late-onset neurodegeneration of retinal ganglion cells (RGCs), optic nerve (ON), and spinal cord motor neurons, preceded by a striking decrease of axonal mitochondria. Surprisingly, we discover that OPTN directly interacts with both microtubules and the mitochondrial transport complex TRAK1/KIF5B, stabilizing them for proper anterograde axonal mitochondrial transport, in a C-terminus dependent manner. Encouragingly, overexpressing OPTN/TRAK1/KIF5B reverses not only OPTN truncation-induced, but also ocular hypertension-induced neurodegeneration, and promotes striking ON regeneration. Therefore, in addition to generating new animal models for NTG and ALS, our results establish OPTN as a novel facilitator of the microtubule-dependent mitochondrial transport necessary for adequate axonal mitochondria delivery, and its loss as the likely molecular mechanism of neurodegeneration.

## Introduction

Axonopathy is a common early feature of central nervous system (CNS) neurodegenerative diseases ^1, 2^, especially in amyotrophic lateral sclerosis (ALS) ^3^ and glaucoma ^4^. ALS patients suffer from progressive neurodegeneration of axons sent out by cortical and spinal cord motor neurons. Similarly, patients with glaucoma, the most common cause of irreversible blindness, undergo degeneration of optic nerve (ON), formed by the unidirectional projection axons sent exclusively from retinal ganglion cells (RGCs), and retrograde RGC death. Although elevated intraocular pressure (IOP) is a risk factor for glaucoma, up to one-third to half of glaucoma patients have normal or even below average IOP, a condition called normal tension glaucoma (NTG) ^5^. Various optineurin (OPTN) mutations are associated with both familial and sporadic ALS ^6^ and NTG ^7^, which establishes that the two CNS axonopathies are pathogenetically related through OPTN-mediated mechanisms. Although OPTN’s role as a selective autophagy receptor in mitophagy has been the focus for pathogenesis studies ^8–11^, how OPTN dysfunction leads to CNS neurodegeneration is still not clear, in part due to the lack of animal neurodegeneration models induced by OPTN mutations.

Mitochondria produce most of the cellular ATP by oxidative phosphorylation, which is essential for neuron growth, survival, function, and regeneration ^12–14^. After biogenesis in the neuronal soma, mitochondria are anterogradely transported into axons to generate sufficient ATP to meet high axonal energy needs and to buffer axonal Ca^2+^. Proper axonal delivery of mitochondria is crucial for maintaining axon integrity; reduced anterograde movement of mitochondria into axons has been found in mouse models of Alzheimer’s disease (AD) ^15–17^, Huntington’s disease (HD) ^18, 19^, ALS ^20–22^ and glaucoma ^23–27^. The axonal anterograde mitochondrial transport machinery includes adaptor protein trafficking kinesin protein 1 (TRAK1), which connects mitochondria through the mitochondria outer membrane molecule Miro1 to the microtubule motor proteins kinesin-1 family (KIF5A-C) ^13, 14, 28, 29^. Interestingly, OPTN is also known to coordinate intracellular vesicular trafficking through multiple binding partners ^9–11^, but, it has not been linked to mitochondria axonal transport.

By leveraging the simplicity of the mouse retina and ON *in vivo* system, in which RGC somata and axons are anatomically grouped and spatially separated, which permits easy access and straightforward interpretation, we performed a compartmentalized *in vivo* analysis of the neuron-autonomous effects of OPTN dysfunction. Here we reveal a previously unknown function of OPTN in tethering the mitochondria transport complex to microtubules and facilitating axonal mitochondria delivery. We expand these mechanistic findings to show that loss of this function is the likely molecular mechanism of OPTN dysfunction-induced neurodegeneration; and that restoring this pathway leads to profound neuronal survival and axon regeneration, suggesting a promising new strategy for providing neuroprotection and axon regeneration to counter CNS neurodegeneration.

## Results

### OPTN C-terminus truncation (OPTNΔC) induces neurodegeneration in RGCs and spinal cord motor neurons

OPTN is highly expressed in mouse and human RGCs ^30, 31^. We examined the autonomous function of OPTN in RGCs by using a OPTN floxed mouse line, in which exon 12 is flanked by loxP sites ^32^. We previously demonstrated that AAV2 preferentially infects RGCs and that mouse γ-synuclein (mSncg) promoter further restricts Cre expression to RGCs ^33^. Intravitreal injection of AAV2-mSncg-Cre removed exon 12 to create a premature termination code that produced in RGCs a 470 amino acid OPTN C-terminus truncation protein (OPTNΔC) without the two ubiquitin-binding domains (UBDs) in the C-terminus of OPTN, the UBD of ABIN proteins and NEMO (UBAN) and the zinc finger (ZF) domain ^11, 34^ (**Fig. 1A,B**). Multiple mutations in these UBDs are associated with ALS, NTG and juvenile open-angle glaucoma ^34^, including the C-terminus truncation mutation K440Nf*8 and 359fs* in fALS ^35, 36^ and the frame shift mutation D128Rfs*22 with the loss of a much larger C-terminus region found in both NTG ^7^ and ALS ^37^ patients. We then used optical coherence tomography (OCT) to monitor retinal ganglion cell complex (GCC) thickness in living mice, including retinal nerve fiber layer (RNFL), ganglion cell layer (GCL), and inner plexiform layer (IPL), as an *in vivo* indicator of RGC/ON degeneration ^38–43^. There was significant and progressive thinning of the GCC in OPTNΔC eyes compared to contralateral control eyes injected with control AAVs, from 6-8-weeks post AAV-Cre injection (6-8wpi) (**Fig. 1C**). Consistent with these *in vivo* morphological changes, OPTNΔC also caused significant visual function deficits, including decreased amplitude of pattern electroretinogram (PERG), a sensitive electrophysiological assay of general RGC physiological function ^42–45^, and decreased visual acuity measured by optokinetic tracking response (OKR) ^42, 43, 46, 47^ at 8wpi (**Fig. 1D,E**). These glaucomatous phenotypes were not associated with IOP elevation in these mice (**Fig. 1F**), consistent with NTG pathogenesis in human patients. Post-mortem histological analysis of retina wholemounts and ON cross-sections confirmed degeneration of RGC somata and axons (**Fig. 1G**): significant neurodegeneration began between 2-4wpi and worsened from 4 to 8wpi.

**Figure 1.**
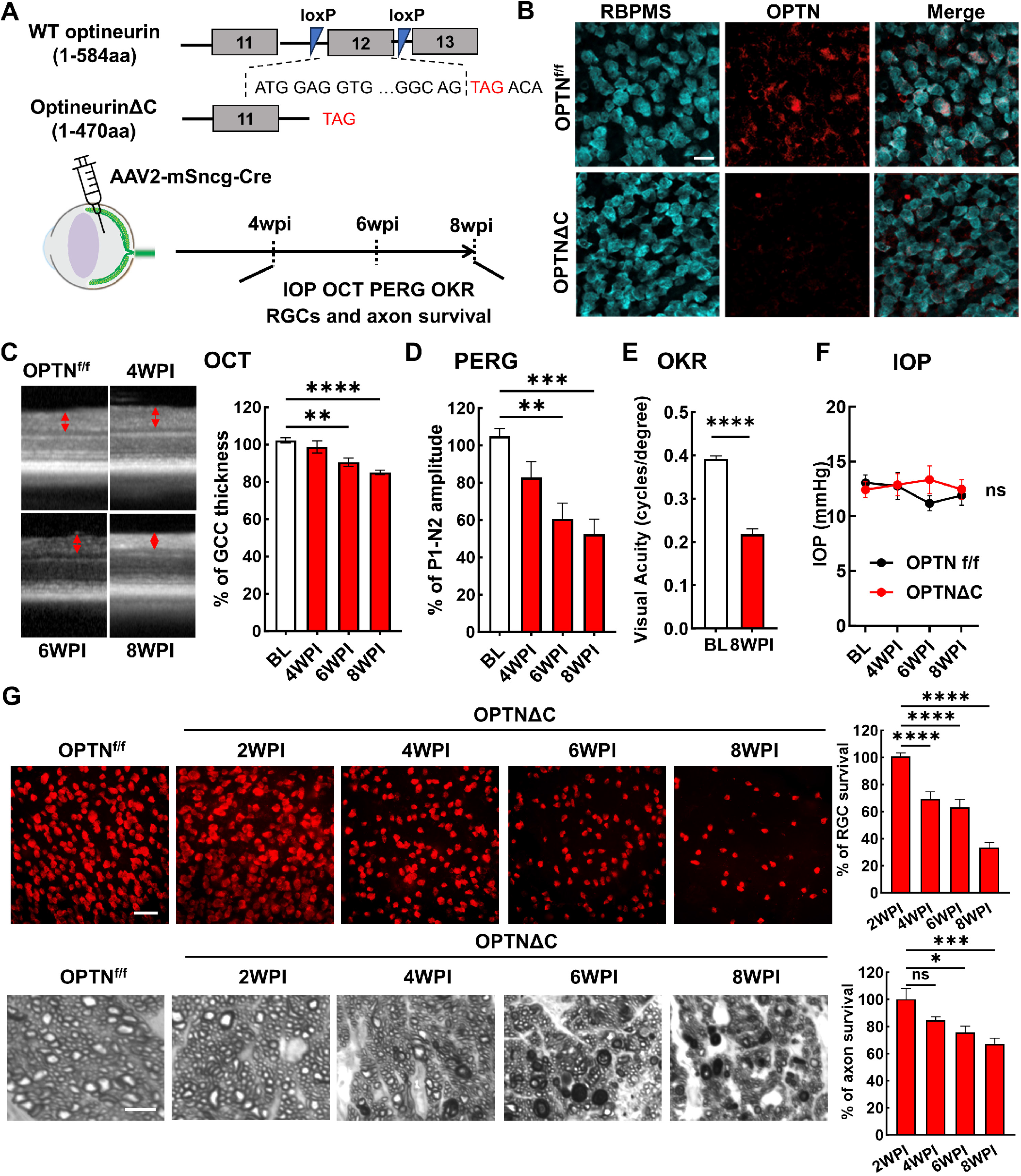
RGC-specific OPTN C-terminus truncation leads to progressive RGC and ON degeneration. A, Exon 12 of the endogenous OPTN gene is flanked by loxP sites in the OPTN^f/f^ mouse line; excision by Cre produces a C-terminus truncated OPTN protein (OPTNΔC). AAV2-mSncg-Cre was intravitreally injected to truncate OPTN specifically in RGCs. *In vivo* measurements of OCT, PERG, OKR, and IOP and histological quantification of surviving RGC somata and axons were performed at 4- 8 weeks post injection (4-8wpi)**. B,** Representative images of retinal wholemounts labeled with RGC marker RBPMS and C-terminus OPTN antibodies. Scale bar, 20 μm. **C,** Representative *in vivo* OCT images of mouse retinas at baseline before AAV-Cre injection, and at 4-8wpi. GCC: ganglion cell complex, including RNFL, GCL and IPL layers; indicated as double end arrows. Quantification is represented as percentage of GCC thickness in the OPTNΔC eyes compared to the contralateral control (CL) eyes. *n* = 10-12 mice. **D,** Quantification of P1-N2 amplitude of PERG at different time points, represented as a percentage of OPTNΔC eyes compared to the CL eyes. *n* = 7-14 mice. **E,** Visual acuity of OPTNΔC eyes and CL eyes measured by OKR at 8wpi. *n* = 7 mice. **F,** IOP of OPTNΔC eyes and CL eyes. *n* = 11 mice. **G**, Upper panel, representative confocal images of retinal wholemounts showing surviving RBPMS-positive RGCs at different time points, Scale bar, 50 µm. Lower panel, light microscope images of semi-thin transverse sections of ON with PPD staining at different time points. Scale bar, 5 µm. Quantification of surviving RGC somata in peripheral retinas and surviving axons in ONs, represented as percentage of OPTNΔC eyes compared to the CL eyes. *n* = 5-12 mice. All the quantification data are presented as means ± s.e.m, *: p<0.05, **: p<0.01, ***: p<0.001, ****: p<0.0001, ns: no significance. **C, D, G** with one-way ANOVA with Dunnett’s multiple comparisons test; **E** with paired Student’s t-test; **F** with two-way ANOVA.

Vglut2 is a pan-RGC marker in the retina and the Vglut2-ires-Cre mouse line ^48^ has been used as a RGC-specific Cre mouse line ^49, 50^. In addition to AAV2-mSncg-Cre delivery, we crossed the Vglut2- ires-Cre line with the OPTN^f/f^ line to generate a transgenic mouse line (OPTN^f/f^::Vglut2-Cre) in which OPTNΔC is expressed only in glutamatergic Vglut2^+^-neurons, including RGCs and spinal cord motor neurons ^51–56^. We confirmed the significant GCC thinning at 8 and 12 weeks of age in this mouse line by *in vivo* OCT imaging (**Fig. 2A**) and visual acuity deficits from 6 to 12 weeks old (**Fig. 2B**). Post-mortem histological analysis of retina wholemounts and ON cross-sections confirmed significant degeneration of RGC somata and axons when they were 12 weeks old but not 4 weeks old (**Fig. 2C,D**). These results show conclusively that RGC-intrinsic C-terminus truncation of OPTN causes autonomous degeneration of RGC somata and axons and visual function deficits without IOP elevation, and therefore establish a clinically relevant mouse NTG-like model with definitive glaucomatous neurodegeneration.

**Figure 2.**
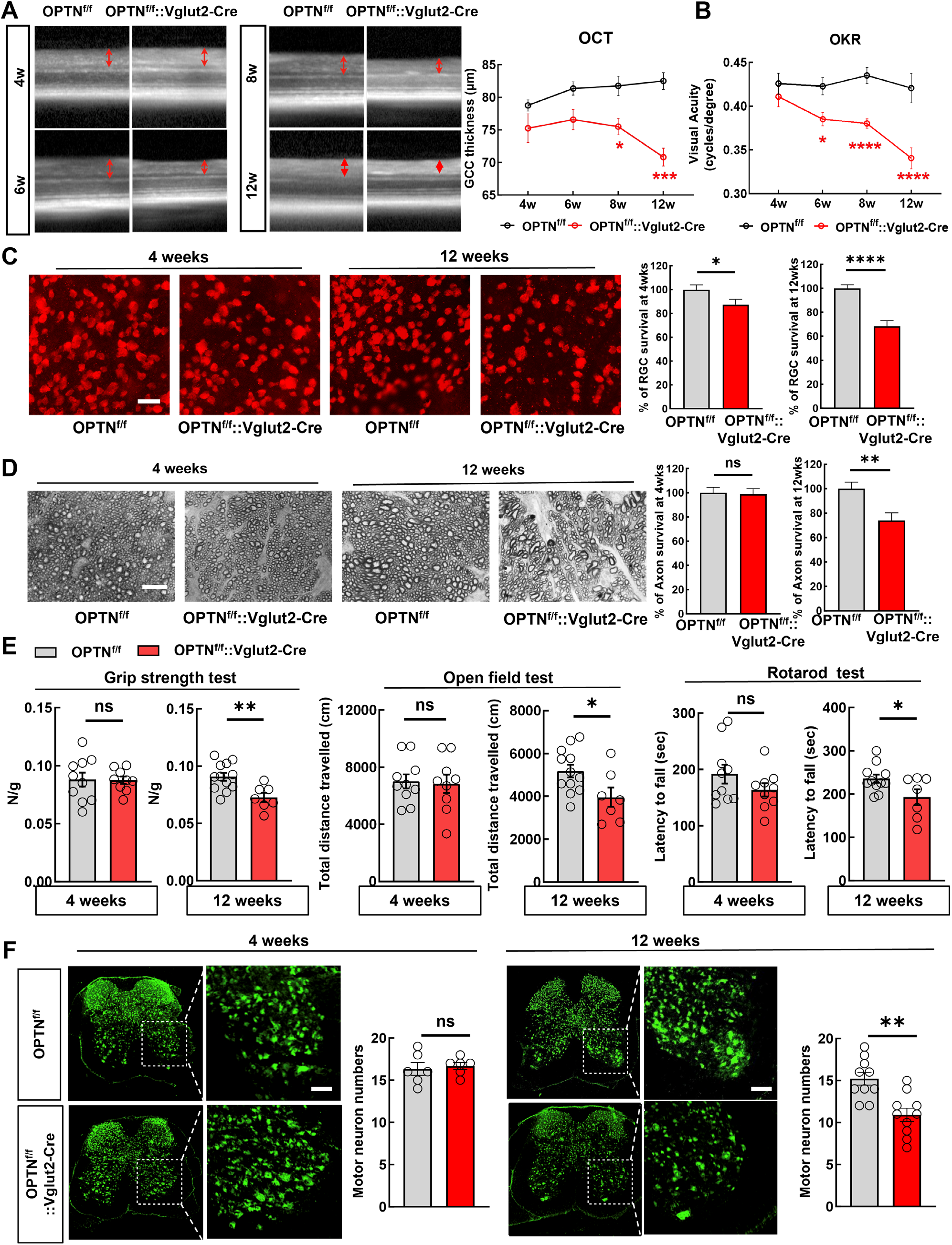
**Vglut2-Cre mediated OPTNΔC causes RGC and spinal cord motor neuron degeneration and ALS-like locomotor deficits**. **A,** Left, representative *in vivo* OCT images of retinas from 4-week-old (4w) to 12w OPTN^f/f^ naïve mice and OPTN^f/f^::Vglut2-Cre mice. Right, quantification of GCC thickness in OPTN^f/f^ naïve and OPTN^f/f^::Vglut2-Cre mouse eyes. *n* = 2-7 mice. **B**, Visual acuity measured by OKR in 4w and 12w OPTN^f/f^ naïve and OPTN^f/f^::Vglut2-Cre mouse eyes. *n* = 3-8 mice. Data are presented as means ± s.e.m, *: p<0.05, ***: p<0.001, ****: p<0.0001, Two-way ANOVA with Sidak’s multiple comparisons test. **C**, Left, representative confocal images of retina wholemounts showing surviving RBPMS-positive (red) RGCs at 4w and 12w. Scale bars, 20 μm; Right, quantification of surviving RGC somata, represented as percentage of OPTN^f/f^::Vglut2-Cre eyes compared with OPTN^f/f^ eyes. **D,** Left, light microscopic images of semi-thin transverse sections of ONs with PPD staining at 4w and 12w. Scale bars, 10 μm. Right, quantification of surviving axons in ONs at 4w and 12w, represented as percentage of OPTN^f/f^::Vglut2-Cre eyes compared with OPTN^f/f^ eyes. n = 8-15 mice. **E**, Behavioral tests of locomotion, including four-paw grip strength, distance traveled in open field test, and latency to fall in rotarod test, were performed in 4- (male = 4, female =5) and 12-weeks old (male = 6, female=1) OPTN^f/f^:: Vglut2-Cre mice and compared to same-age naïve OPTN^f/f^ mice (4 weeks male = 6, female = 4; 12 weeks male = 7 or 6, female = 5). **F**, Immunofluorescent labeling of neurons with NeuN (green) in lumbar segments 1-3 spinal cord sections of the OPTN^f/f^ ::Vglut2-Cre mice and same-age naïve OPTN^f/f^ mice. The motor neurons in the ventral horns were quantified as NeuN^+^ and larger than 500 μm^2^. Scale bars, 100 μm. Quantification of motor neuron survival at 4- or 12-weeks-old are shown to the right. *n* = 5-12 mice. All the quantification data are presented as means ± s.e.m, *: p<0.05, **: p<0.01, ****: p<0.0001, ns: no significance, **C-F** with unpaired t-test.

Surprisingly, we initially found that their body weight of the OPTN^f/f^::Vglut2-Cre mouse line was increased at 12 weeks old (**Fig. S1A**), and then that they showed significant ALS-like locomotor deficits when they were 12 weeks old but not 4 weeks old (**Fig. 2E**), indicating motor neuron degeneration. Indeed, Vglut2-Cre drives YFP expression in spinal cord neurons in both dorsal and ventral horns (**Fig. S1B**). Quantification of large motor neurons in the spinal ventral horns consistently revealed significant motor neuron loss in 12-week-old but not 4-week-old OPTN^f/f^::Vglut2-Cre mice (**Fig. 2F and Fig. S1C**). Taken all together, our results indicate that the C-terminus of OPTN is critical for OPTN’s function; loss in RGCs or spinal motor neurons generates late onset NTG- or ALS-like neurodegeneration due to OPTN dysfunction.

### Dramatic decrease of axonal mitochondria in OPTNΔC-ONs precedes neurodegeneration

OPTN plays an important role in selective autophagy, especially mitophagy, by targeting ubiquitinated mitochondria to autophagosomes ^8, 34, 57, 58^. The OPTN^E478G^ mutant found in ALS patients loses the interaction between the OPTN UBAN domain and ubiquitinated cargos, which disrupts mitophagy and causes cell death in cultured neurons ^59^. The loss of UBDs in the OPTNΔC protein may impair OPTN-mediated neuronal mitophagy, which has been proposed as a potential mechanism for neurodegeneration in glaucoma and ALS ^6, 9, 34^. To examine mitochondria turnover, we used AAV-mediated MitoTimer expression to differentially label young and aged mitochondria in RGCs. MitoTimer is a fusion protein containing a mitochondrial-targeting sequence (MTS) in the time sensitive fluorescence Timer protein that labels newly synthesized young mitochondria green but turns to red when the mitochondria age ^60, 61^. The red/green ratio in OPTNΔC-RGC somata and axons did not differ significantly from that in floxed naïve RGCs at 2wpi, before significant neurodegeneration (**Fig. S2A**), suggesting that RGC mitophagy was not significantly affected by OPTNΔC. Surprisingly, however, the most obvious deficit was a dramatic decrease of mitochondria density in OPTNΔC-ONs (**Fig. S2A**), suggesting that OPTNΔC significantly blocked mitochondrial translocation to axons. To confirm this striking phenotype, we used three complementary approaches to definitively demonstrate a significant decrease in axonal mitochondria in OPTNΔC eyes at 2wpi, before significant neurodegeneration: 1) AAV-mediated RGC expression of another mitochondria tracker containing 4 copies of MTS fused with Scarlet fluorescent protein ^62^ (**Fig. 3A**); 2) Intravitreal injection of cell-permeant and fixable mitochondrion-selective dye, MitoTracker Orange CMTMRos ^63–65^, to label healthy mitochondria with intact mitochondrial membrane potential (**Fig. 3B,C**); 3) Transmission electron microscope (TEM) quantification of axonal mitochondria in ON cross-sections (**Fig. 3D**). All three methods showed dramatically decreased total or healthy mitochondria in RGC axons with OPTNΔC. In contrast, there was no significant difference in mitochondria labeling in RGC somata (**Fig. S2B,C**), and no obvious morphological changes of axonal mitochondria in ONs (**Fig. S2D**). OPTNΔC did not affect general passive axonal transport of cytosol fluorescence protein or active axonal transport of the anterograde axonal tracer cholera toxin subunit B ^66, 67^ (**Fig. S2E,F**). Therefore, OPTNΔC causes an axon-specific mitochondrial transport deficit that precedes significant neurodegeneration.

**Figure 3.**
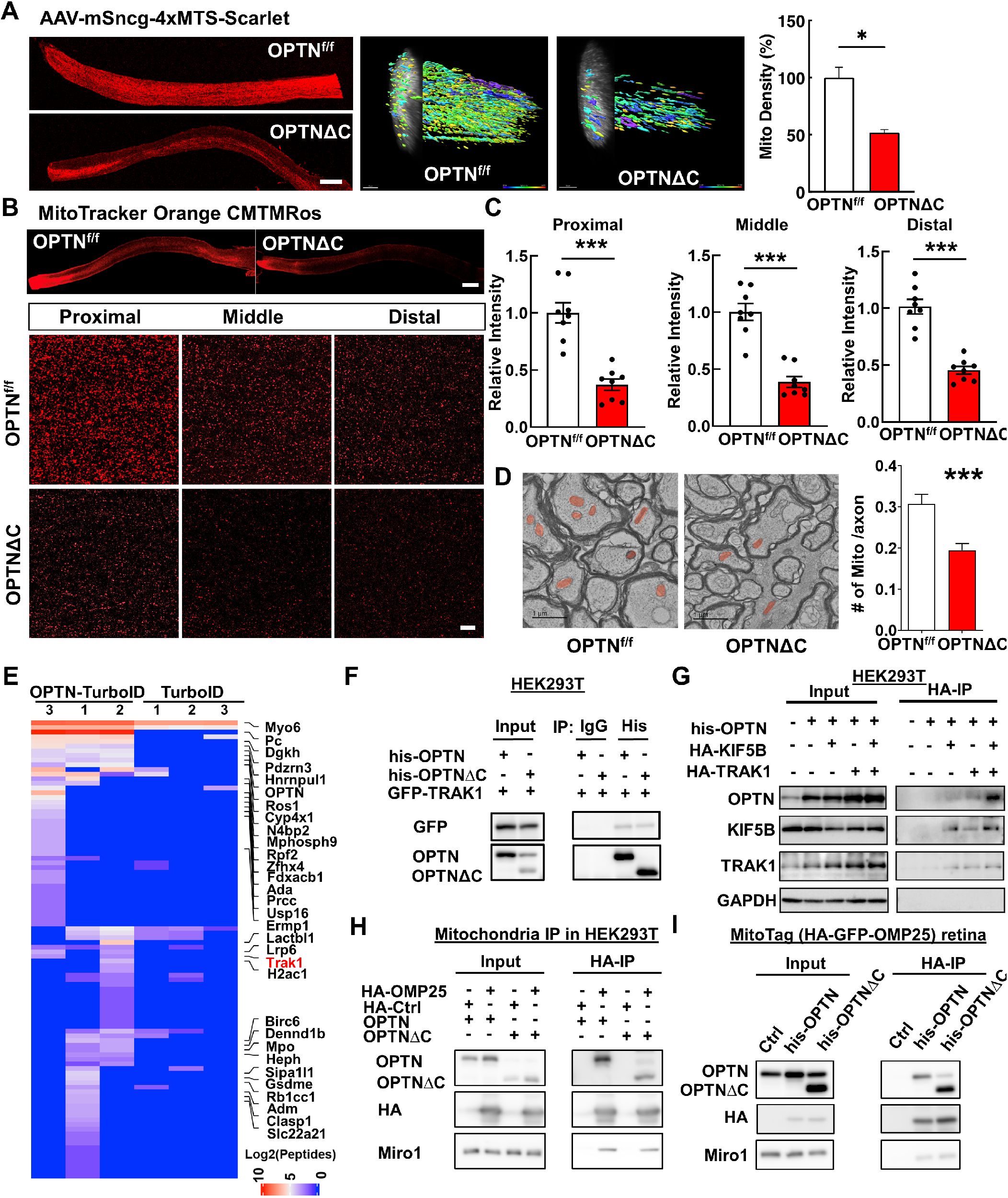
Dramatic decrease of axonal mitochondria in OPTNΔC-ONs precedes neurodegeneration; OPTN directly interacts with the TRAK1-KIF5B-mitochondria transport complex. A, Left, representative images of ON longitudinal sections 2 weeks after intravitreal injection of AAV-4xMTS- Scarlet or AAV-Cre + AAV-4xMTS-Scarlet in OPTN^f/f^ mice. Scale bar, 200 μm. Middle, 3-dimensional (3D) reconstruction of axon mitochondria in ONs showing mitochondrial density. Mitochondrial sphericity is shown in the color bar. Right, quantification of mitochondrial density, represented as a percentage of OPTNΔC eyes compared to the CL eyes. *n* = 5 mice. **B,** Representative images of ON wholemount labeled by MitoTracker Orange CMTMRos 2 weeks after intravitreal injection of AAV-Cre. Scale bar, 200 μm. Higher magnification images of ON segments with labeled mitochondria are shown at the bottom. **C,** Quantification of mitochondrial density of proximal, middle and distal ON wholemounts, represented as a percentage of OPTNΔC eyes compared to the CL eyes. *n* = 5 mice. **D,** Representative TEM images of ON cross-sections (10,000 x) 2 weeks after intravitreal injection of AAV-Cre. Mitochondria are labeled in pseudo color red. Quantification of the mitochondria numbers per axon in ONs. *n* = 4 mice. All the quantification data are presented as means ± s.e.m, *: p<0.05, ***: p<0.001, paired Student’s t-test. **E,** Heatmap of enriched OPTN-interacting proteins in RGCs identified by *in vivo* TurboID and compared by OPTN-TurboID *vs* TurboID alone. **F**, Co-IP analysis of HEK293T cells with corresponding overexpression. α-his antibodies were used to IP OPTN and corresponding antibodies for recognizing individual proteins. **G,** Co-IP analysis of HEK293T cells with corresponding overexpression. α-HA magnetic beads were used for IP KIF5B or TRAK1 and corresponding antibodies of individual proteins for recognition. **H**, Co-IP analysis of HEK293T cells with corresponding overexpression. α-HA magnetic beads were used for IP HA-GFP-OMP25-labeled mitochondria and corresponding antibodies of individual proteins for recognition. **I,** Co-IP analysis of MitoTag mouse retinas with corresponding overexpression. α-HA magnetic beads were used for IP HA-GFP-OMP25-labeled mitochondria and corresponding antibodies of individual proteins for recognition.

### OPTN directly interacts with the TRAK1-KIF5B-mitochondria transport complex

OPTN serves multiple functions through direct interaction with various signaling, adaptor, and motor molecules ^9^. Our results indicate a novel role of OPTN in axonal mitochondria transport, but none of its known binding partners are involved in this process. We therefore employed a proximity-dependent biotin identification assay, TurboID ^68, 69^, to systematically profile OPTN-associated proteins in RGCs *in vivo.* We first confirmed the feasibility of this strategy using OPTN fused with the mutated *E. coli* biotin ligase (TurboID) in cultured HEK293 cells (**Fig. S3A**), and then optimized the mouse *in vivo* biotinylation conditions after AAV-mediated expression of OPTN-TurboID in RGCs (**Fig. S3B-D**). OPTN-TurboID catalyzed the biotinylation of proteins even just transiently and spatially proximal to OPTN in the natural environment of RGCs. These proteins were then purified by streptavidin-conjugated beads from whole retina lysate without the need for retinal cell dissociation and RGC isolation and identified using liquid chromatography and mass spectrometry (LC-MS). The top 30 enriched proteins in OPTN-TurboID-RGCs compared to control RGCs expressing TurboID alone are listed in the heatmap (**Fig. 3E**). The enriched proteins included OPTN itself, another well-known OPTN-interacting protein Myo6 ^34^, and four OPTN-interacting proteins (Pc, Hnrnpul1, Rb1cc1, and Clasp1) that were recently identified by a different *in vitro* proximity assay with OPTN ^70^. Most interestingly, our *in vivo* RGC proximity labeling assay identified a previously unknown OPTN-interacting protein, TRAK1, a crucial adaptor protein that attaches mitochondria through Miro1 to the microtubule-based molecular motor KIF5B for anterograde axonal transport of mitochondria ^14^. To confirm the proximity labeling result, we used co-immunoprecipitation in HEK293 cells to demonstrate that GFP-TRAK1 can be pulled down by OPTN immunoprecipitation (**Fig. 3F**). OPTN also can be pulled down by HA-tagged TRAK1 or KIF5B immunoprecipitation, and more intriguingly, significantly more OPTN was pulled down when both KIF5B and TRAK1 were overexpressed (**Fig. 3G**), indicating the strong interaction of OPTN with the mitochondria transport complex TRAK1/KIF5B. To further confirm that OPTN is associated with mitochondria, we performed mitochondria immunoprecipitation with HA-tagged mitochondria membrane protein OMP25 in HEK293 cells. The HA-mediated mitochondria pulldown co-immunoprecipitated with OPTN (**Fig. 3H**). This mitochondria-OPTN co-immunoprecipitation was also confirmed in retinas of the MitoTag mice *in vivo* (**Fig. 3I**). To our surprise, however, OPTNΔC also co-immunoprecipitated with TRAK1 and mitochondria (**Fig. 3F,H,I**), suggesting that the C-terminus truncation of OPTN does not affect the direct binding of OPTN with the TRAK1-KIF5B-mitochondria transport complex and that the neurodegeneration associated with OPTNΔC is not due to the loss of this direct interaction. In summary, we revealed a major axonal mitochondria transport deficit in the OPTNΔC-neurons that precedes neurodegeneration and a previously unknown interaction between OPTN and the mitochondria transport complex, including TRAK1, KIF5B and mitochondria. These findings suggest that OPTN plays a direct and important role in axonal transport of mitochondria. They cannot explain how OPTNΔC affects axonal mitochondria delivery, however, as the loss of the C-terminus does not affect the direct interaction between OPTN and the mitochondria transport complex.

### OPTN tethers the KIF5B-TRAK1 complex to microtubules in a C-terminus dependent manner to deliver adequate numbers of axonal mitochondria

We previously demonstrated that TRAK1 activates KIF5B-mediated mitochondria transport along microtubules, using a well-controlled *in vitro* reconstitution motility assay with recombinant TRAK1 and KIF5B proteins and immobilized microtubules on coverslips^71^. Here we used the analogous assay to investigate the role of OPTN in the motility of the KIF5B-TRAK1 transport complex on microtubules. Using purified mNeonGreen tagged OPTN (mNG-OPTN) and OPTNΔC (**Fig. S4A**), we found, surprisingly, that OPTN itself bound directly to microtubules (**Fig. 4A**), with rapid binding and unbinding kinetics (**Movie S1**). In contrast, OPTNΔC lost the microtubule-binding ability (**Fig. 4A, Movie S1**), indicating that OPTN interacts with microtubules directly in a C-terminus dependent manner. Moreover, by adding lysates of cells overexpressing mNG-OPTN to surface immobilized microtubules, we confirmed that OPTN interacts with microtubules in the presence of other cellular components (**Fig. 4B, Movie S2**). To further confirm the microtubule-bound OPTN in neurons, we expressed EGFP-tagged OPTN or OPTNΔC in cultured primary mouse hippocampal neurons and co-labeled with a live cell microtubule dye SPY555-tubulin. Consistently, super-resolution imaging revealed microtubule-bound full length OPTN in hippocampal neurons, in dramatic contrast to the obviously dispersed pattern of OPTNΔC (**Fig. 4C**). Moreover, AlphaFold2 predicts the direct interaction between a portion of the C-terminus of OPTN and alpha-tubulin (**Fig. 4D**), further confirmed the C-terminus dependent OPTN binding to microtubules.

**Figure 4.**
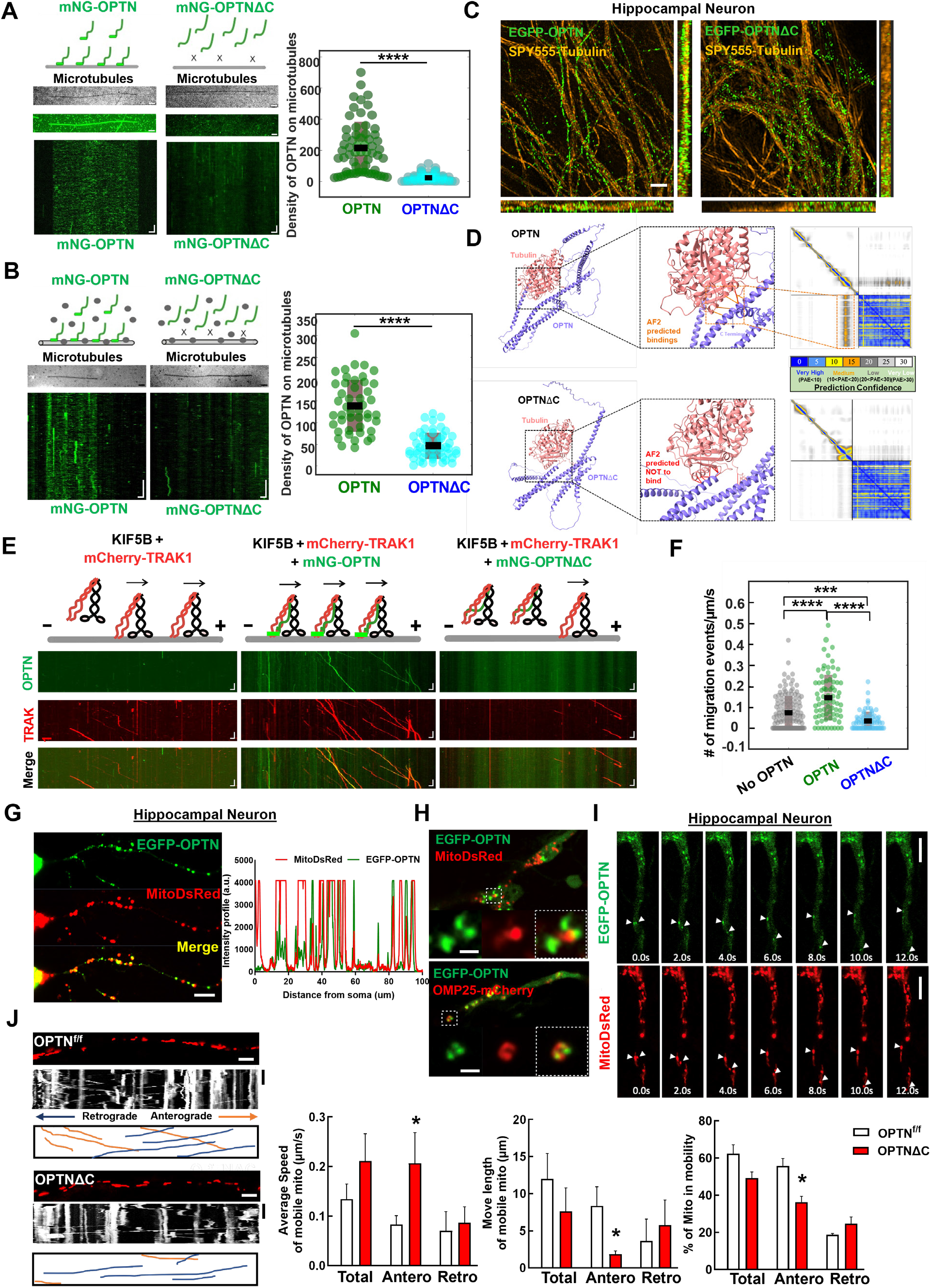
**OPTN tethers KIF5B-TRAK1 complex to microtubules in a C-terminus dependent manner for adequate axonal mitochondria delivery**. **A,** *In vitro* reconstitution assay: OPTN on immobilized microtubules. Left (top to bottom): Schematic representing interaction of OPTN/OPTNΔC with microtubules, IRM image of microtubules, maximum intensity projection, and kymograph of 20 nM mNG-OPTN or 0.1 µM mNG-OPTNΔC. Horizontal scale bar = 2.1 μm, vertical scale bar, 4 seconds. Right, quantification of density of OPTN/OPTNΔC on microtubules. *n* = 3 experiments. ****: p <0.0001, t-test. **B,** Lysates of mNG-OPTN or mNG-OPTNΔC overexpressing cells on immobilized microtubules. Left (top to bottom): Schematic representing interaction of cell lysates expressing OPTN/OPTNΔC with microtubules, IRM image of microtubules, and kymograph of mNG-OPTN or mNG-OPTNΔC. Horizontal scale bar, 2.0 μm, Vertical scale bar, 10 seconds. *n* = 3 experiments. Right, quantification of density of OPTN/OPTNΔC on microtubules. n = 3 experiments, ****: p <0.0001, t-test. **C,** SIM super-resolution images of cultured mouse E15 hippocampus neuron transfected with EGFP-OPTN or EGFP- OPTNΔC, and stained for microtubules with SPY555-tubulin. Scale bars, 2 μm. **D,** The AlphaFold2 predicated interaction between OPTN or OPTNΔC and Tubulin alpha-1A. The prediction confidence is visualized as heatmap of Predicted Aligned Error (PAE) plots. The x-axis and y-axis of the plot represent the sequence of amino acids in the two proteins. Each dot is color-coded in PAE in the grid, corresponding to the pair of amino acids in both proteins. **E,** *In vitro* reconstitution motility assay of immobilized microtubules. (top to bottom) Schematic representation of KIF5B-TRAK1 transportation complex with or without OPTN or OPTNΔC on microtubules, kymograph of mNG-OPTN or mNG-OPTNΔC and mCherry-TRAK1 walking to plus end of microtubules in the presence of unlabeled KIF5B. Horizontal scale bar, 2 μm; vertical scale bar, 10 seconds. **F,** Frequency of migration events (/μm/s) of complexes of KIF5B-TRAK1 (n = 133), KIF5B-TRAK1-OPTN (n = 73), KIF5B-TRAK1-OPTNΔC (n = 71), *n* = 3 experiments. ****: *p* < 10^-7^, ***: *p* =0.0002, t-test. **G**, Representative confocal images of cultured hippocampal neuron axons co-transfected with meGFP-OPTN with MitoDsRed to show the colocalization of OPTN and mitochondria in axons. The intensity profile analysis is shown in the right panel. Scale bar, 20 μm**. H**, Representative confocal images of cultured hippocampal neuron axons co-transfected with meGFP-OPTN with MitoDsRed or OMP25-mCherry. Higher magnification images of mitochondria are shown in the right lower panels. Scale bar, 5 μm**. I**, Time-lapse images of cultured hippocampal neuron axons co-transfected with meGFP-OPTN with MitoDsRed. White arrow heads indicate the colocalized meGFP-OPTN and mitochondria that are moving together in axons. Scale bar, 20 μm**. J,** Mitochondrial movement (anterograde movement: orange, retrograde movement: blue) in OPTN^f/f^ or OPTNΔC hippocampal neuron axons transfected with MitoDsRed. The first frame (time = 0s) of live imaging series is shown with the kymograph. Quantifications of average speed of mobile mitochondria, move length of mobile mitochondria and percentage of mitochondria in mobility are shown to the right. n = 12-15 mitochondria from 3 axons per group. Horizontal scale bar, 10 μm, Vertical sale bar, 1 minute. Quantification data are presented as means ± s.e.m, *: p<0.05 with t-test.

We next found that adding purified KIF5B did not affect the binding of OPTN to microtubules, and that OPTN only very rarely migrated along microtubules (**Fig. S4B**), suggesting there is no, or very weak/transient interaction between KIF5B and OPTN in the absence of TRAK1. Similarly, adding TRAK1 did not affect the binding of OPTN to microtubules, and we detected no movement of TRAK1 and OPTN in the absence of motor molecule KIF5B (**Fig. S4C**). We next added a combination of all three purified components to the surface immobilized microtubules. Consistent with our previous finding that TRAK1 activates KIF5B ^71^, we detected migration of KIF5B-TRAK1 towards the microtubule plus-end (**Fig. 4E**, **Movie S3**). Significantly, in the presence of both TRAK1 and KIF5B, OPTN co-migrated with the KIF5B-TRAK1 complex along microtubules, and markedly increased the frequency of migration events of the tripartite complex on microtubules (**Fig. 4E,F**, **Movie S3**). We attributed these changes in motility to the direct interaction of OPTN with the microtubule surface, which, by tethering and stabilizing the KIF5B-TRAK1 transport complex with the microtubule, produces longer run times/run lengths at reduced velocities (**Fig. S5A,B**). To test this hypothesis, we repeated the experiment with the same concentrations of KIF5B-TRAK1 and OPTNΔC, removing the ability of OPTN to bind microtubules (**Fig. 4E**, **Movie S3**). Consistent with our hypothesis, removing the C-terminus of OPTN decreased the frequency of migration events of the KIF5B-TRAK1 complex (**Fig. 4E.F, Movie S3**), as well as the run length and run time, while increasing the velocity compared to that in the presence of full length OPTN (**Fig. S5A,B**). Importantly, we found that the presence of OPTNΔC reduced the number of migration events even below the number in the absence of OPTN (**Fig. 4F**), suggesting that OPTNΔC not only ceases to function but may also actively prevent the landing of the mitochondria transport complex onto microtubules. Based on our previous result that OPTNΔC continues to bind to TRAK1-KIF5B- mitochondria (**Fig. 3F-I**), but loses its microtubule binding capability (**Fig. 4A-D**), it is plausible that OPTNΔC traps the TRAK1-KIF5B-mitochondria complex and decreases its binding to microtubules, therefore hindering the anterograde axonal transport of mitochondria. It is also worth noting that the measured events without OPTN and with OPTNΔC in **Fig. S5A,B** are essentially the same, because the TRAK1-KIF5B complex on the microtubules is the same; there is no OPTN in either situation because OPTNΔC does not bind to microtubules. Therefore, it is not surprising that the No OPTN group and the OPTNΔC group do not differ in **Fig. S5A,B**.

To validate these *in vitro* results in *in vivo* conditions, we next performed the same motility assay using lysates of cells overexpressing fluorescence tagged OPTN and TRAK1 supplemented with purified KIF5B (**Fig. S5C**). We found that the motility of the KIF5B-TRAK1-OPTN/OPTNΔC complex in cell lysates is qualitatively identical as our results with purified proteins (**Fig. S5C-F, Movie S4**). We then confirmed that OPTN is partially co-localized with axonal mitochondria in cultured primary hippocampal neurons (**Fig. 4G**). Higher resolution images showed that OPTN surrounds mitochondria matrix labeled by MitoDsRed and colocalizes with mitochondria outer membrane labeled by OMP25-mCherry (**Fig. 4H**). Time-lapse imaging confirmed that axonal OPTN migrates together with axonal mitochondria in hippocampal neurons (**Fig. 4I**). We then used live imaging to measure the motility of axonal mitochondria in cultured hippocampal neurons (**Fig. 4J, Movie S5**): as in the *in vitro* motility assays, OPTNΔC significantly increased the speed of axonal anterograde transportation, but decreased the run length and percentage of moving mitochondria (**Fig. 4J**). We concluded this set of experiments by confirming that OPTN is co-localized with mitochondria in mouse ON (**Fig. S6A**), and that there are significantly fewer moving mitochondria *ex vivo* in ONs freshly isolated from OPTNΔC mice than from naïve mice (**Fig. S6B,C** and **Movie S6**).

Taken all together, these findings demonstrate that OPTN can bind directly to the microtubule surface in a C-terminus dependent manner and to the TRAK1-KIF5B-mitochondria complex in a C-terminus independent manner both *in vitro* and *in vivo*. OPTN therefore serves a previously unappreciated function in microtubule-dependent mitochondria trafficking by tethering the TRAK1-KIF5B-mitochondria transport complex onto microtubules, which, by stabilizing the transport complex, promotes long distance mitochondria transport. When the C-terminus of OPTN is lost or dysfunctions, the loading of the TRAK1- KIF5B-mitochondria complex onto microtubules decreases, which produces significant deficits in axonal mitochondria distribution and ultimately neurodegeneration.

### Increasing axonal mitochondria distribution by overexpressing TRAK1/KIF5B rescues OPTNΔC- induced neurodegeneration

We reasoned that increasing the abundance of the TRAK1-KIF5B motor complex might enhance axonal mitochondria delivery and achieve neuroprotection by overcoming the deleterious effects of OPTNΔC.

Therefore, we generated AAV-HA tagged TRAK1 and KIF5B driven by the RGC-specific mSncg promoter and confirmed their overexpression in RGCs *in vivo* (**Fig. S7A**). We co-injected AAV-Cre + AAV-KIF5B, AAV-Cre + AAV-TRAK1, or AAV-Cre + AAV-KIF5B + AAV-TRAK1 into one eye of the OPTN floxed mice and injected AAV-Cre alone into the contralateral eye for comparison. Both KIF5B and TRAK1 individually enhanced ON mitochondria distribution in OPTNΔC eyes, and KIF5B + TRAK1 together promoted the most significant axonal mitochondria translocation (**Fig. 5A**). We next used *in vivo* OCT imaging to demonstrate that TRAK1 alone or combined with KIF5B, but not KIF5B alone, significantly increased the GCC thickness of OPTNΔC eyes (**Fig. 5B**). Histological analysis of postmortem retina wholemounts and ON transverse sections consistently showed that treatment with KIF5B or TRAK1 alone or together strikingly increased RGC soma and axon survival at 8wpi (**Fig. 5C**). In addition to providing morphological protection, increasing axonal mitochondria by TRAK1/KIF5B significantly preserved visual acuity of the OPTNΔC eyes (**Fig. 5D**). These results demonstrated that overexpressing KIF5B/TRAK1 to enhance the axonal mitochondria transport machinery overcomes the detrimental effect of OPTNΔC on axonal mitochondria distribution and achieves significant neuroprotection.

**Figure 5.**
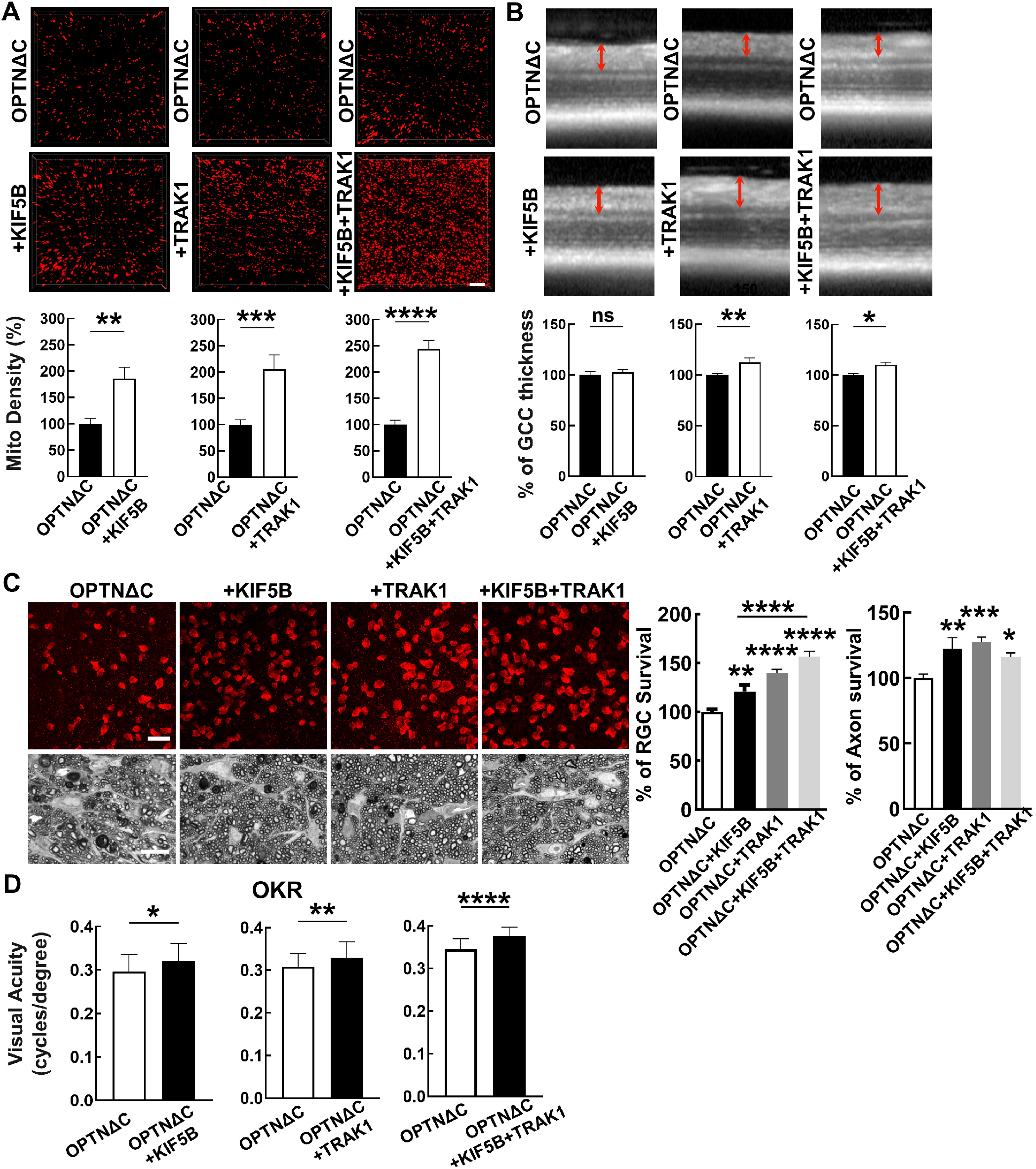
Overexpression of KIF5B and/or TRAK1 rescues the axonal mitochondria deficit and neurodegeneration induced by OPTNΔC. A, Representative images of MitoTracker-labeled ON longitudinal sections. Scale bar, 10 μm. Quantification of Mito Density, represented as a percentage of treated OPTNΔC eyes compared to the CL non-treated OPTNΔC eyes 2 weeks after intravitreal injection of AAV-Cre. *n* = 4-5 mice. Data are presented as means ± s.e.m, **: p<0.01, ***: p<0.001, ****: p<0.0001, paired Student’s t-test. **B,** Representative *in vivo* OCT images of mouse retinas at 8wpi. Quantification is represented as percentage of GCC thickness in the treated OPTNΔC eyes compared to the CL non-treated OPTNΔC eyes. *n* = 7-10 mice. Data are presented as means ± s.e.m, ns, no significance; *: p<0.05, **: p<0.01, paired Student’s t-test. **C,** Upper panel, representative confocal images of retinal wholemounts showing surviving RBPMS-positive RGCs at 8wpi. Scale bar, 20 µm. Lower panel, light microscope images of semi-thin transverse sections of ON with PPD staining at 8wpi. Scale bar, 10 µm. Quantification of surviving RGC somata in peripheral retinas and surviving axons in ONs, represented as percentage of treated OPTNΔC eyes compared to the CL non-treated OPTNΔC eyes. *n* = 8-12 mice. Data are presented as means ± s.e.m, *: p<0.05, **: p<0.01, ***: p<0.001, ****: p<0.0001, one-way ANOVA with Tukey’s multiple comparisons test. **D,** Visual acuity of treated and CL non-treated OPTNΔC eyes measured by OKR at 8wpi. *n* = 8-16 mice. Data are presented as means ± s.e.m, *: p<0.05, **: p<0.01, ****: p<0.0001, paired t-test.

### Ocular hypertension also decreases ON mitochondria transportation; OPTN/TRAK1/KIF5B reverses glaucomatous ON mitochondria deficits and neurodegeneration

Axonal mitochondria transport has also been shown to be deficient in the mouse microbead-induced ocular hypertension glaucoma model ^23, 24, 27^. We next asked whether a similar deficit also occurs in the silicone oil-induced ocular hypertension (SOHU) mouse glaucoma model, which we recently developed to closely mimic human secondary glaucoma ^38–41^. We examined ON mitochondria numbers and motilities at an early time point, before the onset of significant glaucomatous neurodegeneration. Intriguingly, we found that the total number of axonal mitochondria and the speed and mobile time of moving mitochondria were significantly decreased in glaucomatous ONs at 1-week post SO injection (1wpi), whereas the stationary time of axonal mitochondria was significantly increased (**Fig. 6A, Movie S7**). These results suggest that impaired axonal mitochondria transport is a common feature of both NTG and ocular hypertension glaucoma. AAV-mSncg-mediated OPTN/TRAK1/KIF5B overexpression in RGCs reversed the axonal mitochondria deficits by increasing the total number of ON mitochondria and the speed and mobile time of moving mitochondria but decreasing the stationary time of mitochondria (**Fig. 6A, Movie S7**). We then investigated RGC/ON morphology and visual function in the SOHU glaucoma model at 3wpi, when there is significant IOP elevation (**Fig. S7B**). *In vivo* OCT imaging showed significant thinning of the GCC in SOHU eyes compared to contralateral control eyes in control group animals injected with control AAVs at 3wpi (**Fig. 6B**). Consistently, overexpression of OPTN alone or together with TRAK1 and KIF5B, significantly increased GCC thickness in SOHU eyes (**Fig. 6B**). *In vivo* assessment of visual acuity by OKR and of RGC electrophysiology function by PERG demonstrated that increasing axonal mitochondria by OPTN alone or together with TRAK1/KIF5B significantly preserved visual function of the SOHU glaucoma eyes (**Fig. 6C,D**). Histological analysis of postmortem retina wholemounts consistently showed that treatment with OPTN alone or OPTN/TRAK1/KIF5B strikingly increased RGC survival throughout the whole retina and axon survival in the ONs (**Fig. 6E**). These results demonstrated that axonal mitochondria deficits are a common feature of glaucoma with or without IOP elevation and that increasing ON mitochondria delivery by enhancing the axonal mitochondria transport machinery (OPTN/KIF5B/TRAK1) is a promising neuroprotection strategy for glaucomatous neurodegeneration.

**Figure 6.**
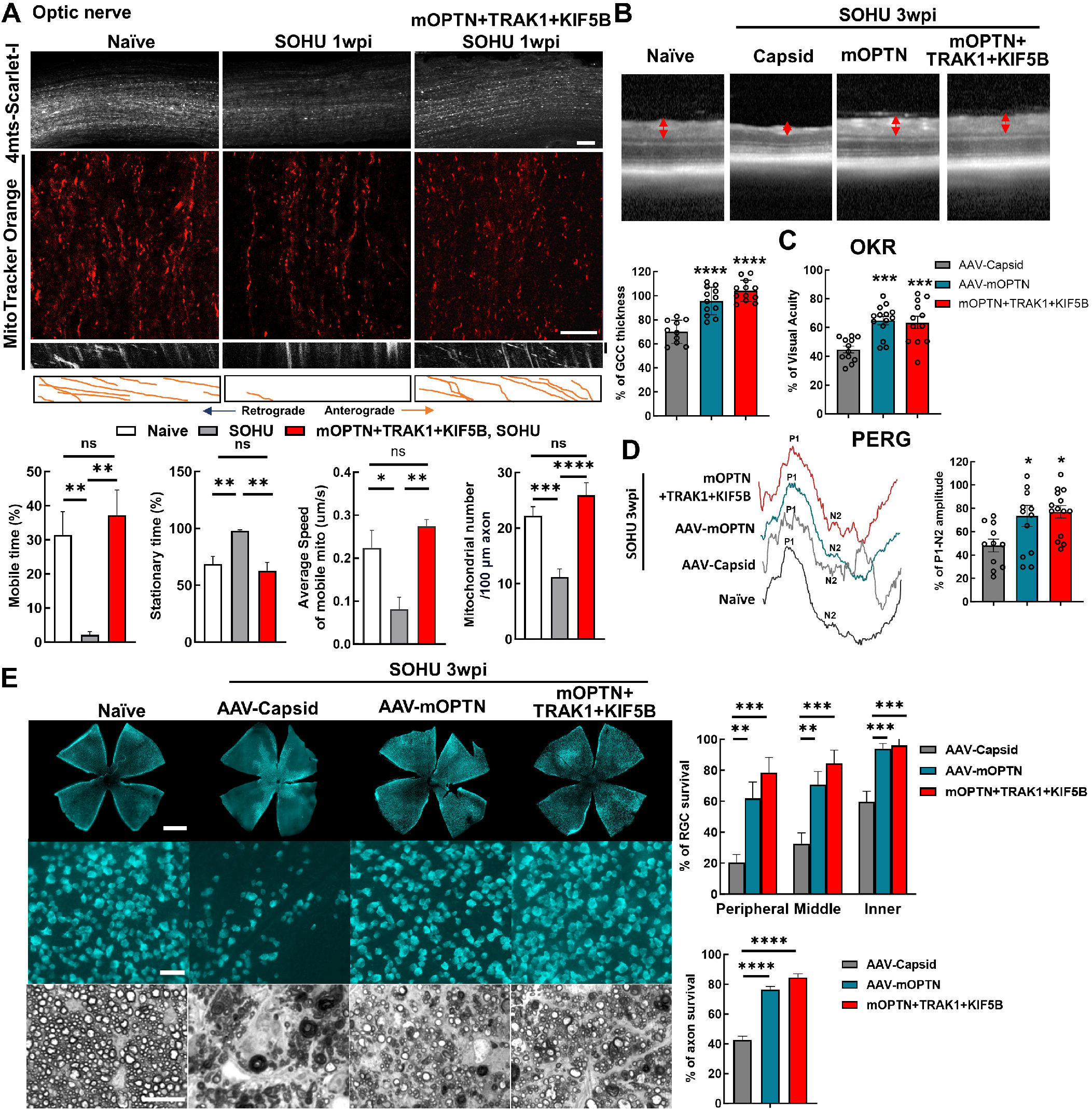
Ocular hypertension decreases ON mitochondrial transportation; OPTN/TRAK1/KIF5B reverses glaucomatous ON mitochondrial deficits and neurodegeneration in SOHU glaucoma model. A, (top to bottom) Representative confocal images of ON wholemounts from naïve mice, SOHU mice at 1wpi and SOHU mice at 1wpi with mOPTN+TRAK1+KIF5B overexpression and AAV-4MTS-Scarlet labeling, Scale bars, 50 μm; representative ON wholemount images of the three groups with MitoTracker Orange labeling, Scale bars, 20 μm; kymograph and traces of MitoTracker labelled mitochondria movement along ON axons in the three groups. Vertical scale bar, 1 minute; quantification of each mitochondria’s time in motion and time stationary, average speed of each mobile mitochondrion and total mitochondria number in the axon. n = 17–30 mitochondria from 3 axons per group. Data are presented as means ± s.e.m, *: p<0.05, **: p<0.01, ***: p<0.001, ****: p<0.0001, ns, no significance, with Student’s t-test. **B**, Representative *in vivo* OCT images of SOHU mouse retinas at 3wpi. Quantification is represented as percentage of GCC thickness of glaucomatous eyes compared to the contralateral control eyes. *n* = 10-12 mice. **C,** Visual acuity measured by OKR at 3wpi, represented as percentage of glaucomatous eyes compared to the contralateral control eyes. *n* = 12-14 mice. **D,** Left: representative waveforms of PERG of SOHU mice at 3wpi. Right: quantification of P1-N2 amplitude of PERG at 3wpi, represented as percentage of glaucomatous eyes compared to the contralateral control eyes. *n* = 11-15 mice. **E,** Left, (top to bottom) representative confocal images of retina wholemounts showing surviving RBPMS-positive (cyan) RGCs at 3wpi. Scale bars, 50 μm; light microscopic images of semi-thin transverse sections of ON with PPD staining at 3wpi. Scale bars, 10 μm. Right, Quantification of surviving RGC somata and axons at 3wpi, represented as percentage of glaucomatous eyes compared with the sham contralateral control eyes. n = 10 mice. (**B-E**) All the data are presented as means ± s.e.m, *: p<0.05, **: p<0.01, ***: p<0.001, ****: p<0.0001, one-way ANOWA with Turkey’s multiple comparison tests, compared to AAV-Capsid treated control group.

### The OPTN/KIF5B/TRAK1 complex promotes striking ON regeneration after ON crush (ONC)

Mitochondria axonal targeting has also been linked to CNS axon regeneration ^64, 65, 72–75^. We reasoned that the highly efficient mitochondria transport complex OPTN/KIF5B/TRAK1 may also promote ON regeneration. To test this hypothesis, we overexpressed these genes in RGCs and tested whether they increased axon regeneration and RGC survival after ONC injury. Excitingly, OPTN/KIF5B/TRAK1 promoted striking ON regeneration comparable to that induced by PTEN deletion (**Fig. 7A**); traumatic RGC survival was also increased (**Fig. S7C**). Parallel molecular mechanisms downstream of OPTN/KIF5B/TRAK1 and PTEN seem to be involved in axon regeneration because the PTEN deletion-induced axon regeneration was not affected by OPTN truncation or KIF5B deletion (**Fig. 7A**). On the other hand, overexpression of OPTN/KIF5B/TRAK1 in the PTEN KO mice generated much more potent ON regeneration than either alone (**Fig. 7A**). Some regenerating axons extended to or even traversed the optic chiasma at 4wpc (**Fig. 7B**), indicating an additive or synergistic effect.

**Figure 7.**
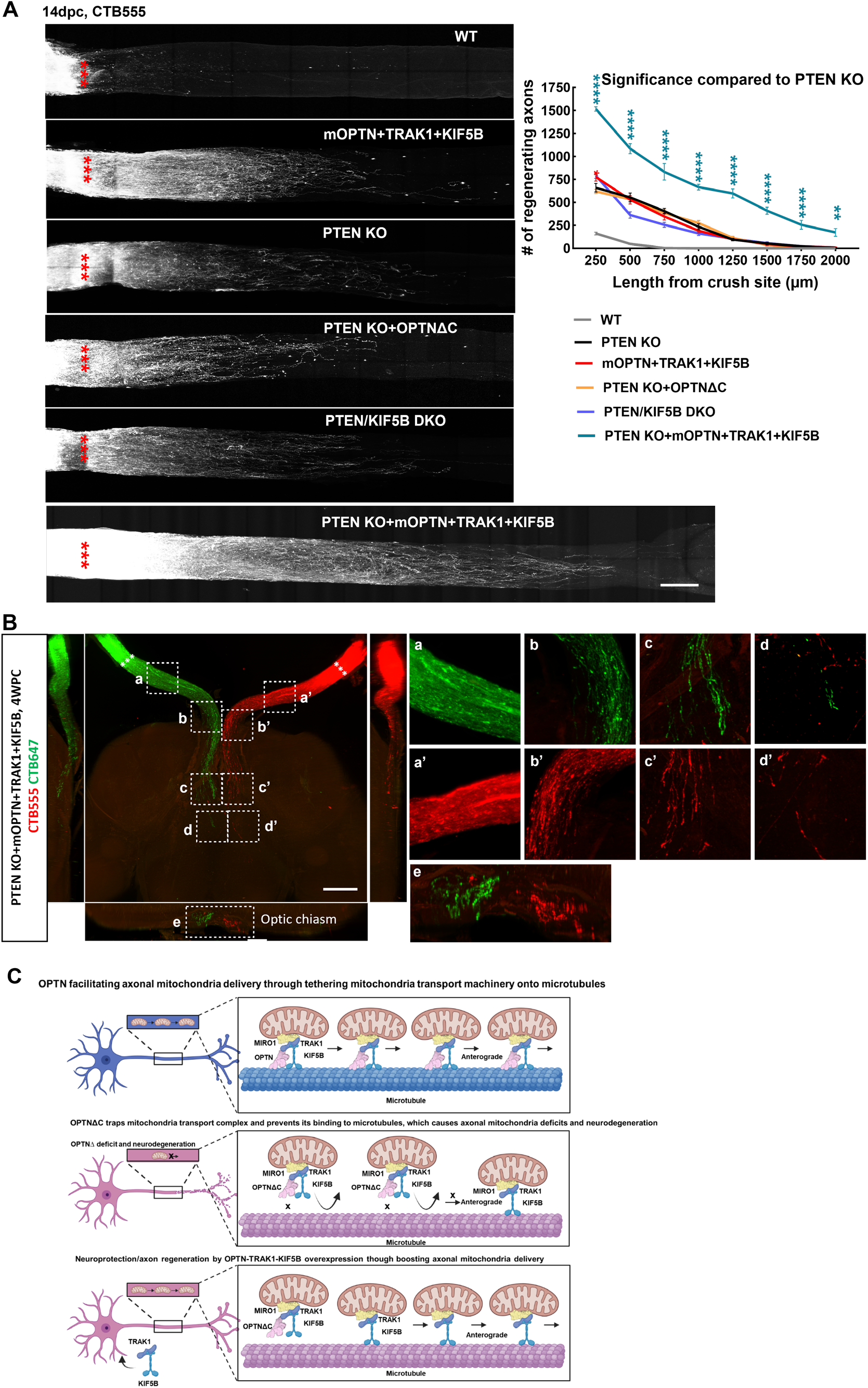
**The OPTN/KIF5B/TRAK1 complex promotes striking ON regeneration after ONC. A**, Left, confocal images of ON wholemounts after optical clearance showing maximum intensity projection of regenerating axons labeled with CTB-Alexa 555 at 14dpc. Scale bar, 250 μm. ***: crush site. Right, quantification of regenerating axons at different distances distal to the lesion site. n = 4-9. Data are presented as means ± s.e.m, *: p<0.05, **: p<0.01, ****: p<0.0001, two-way ANOVA with Dunnett’s multiple comparisons test, compared to PTEN KO group. **B**, Light-sheet fluorescent images (bottom view, sagittal view and coronal view) of regenerating axons in ONs, optic chiasm, and optic tract in PTEN KO mice with mOPTN+TRAK1+KIF5B overexpression at 4 weeks post ONC (4wpc). Regenerating axons were labeled with CTB-Alexa 555 and CTB-Alexa 647 in both eyes separately. Higher magnification images of framed regions (a-e, a’-e’) are shown to the right. Scale bar, 500μm. **C,** Models of OPTN physiological role in axonal mitochondria transport, dysfunctional OPTNΔC in jeopardizing axonal mitochondria distribution and inducing neurodegeneration, and neuroprotection and axon regeneration of OPTN-TRAK1-KIF5B by increasing axonal mitochondria delivery.

## Discussion

OPTN is involved in diverse cellular functions through its broad binding partner network, including inhibition of NF-κB and osteoclast differentiation, maintenance of the Golgi apparatus, coordination of intracellular vesicle trafficking, and autophagy. Many of these processes have been linked to neurodegeneration, especially mitophagy ^9, 34^. However, a recent study found that OPTN-mediated mitophagy is restricted to neuronal somata and is scarcely detectable in axons, and that the OPTN mutation associated with ALS does not affect mitochondria content in neuronal somata ^59^. Mitophagy also remains normal in OPTN mutations associated with glaucoma ^76^. Therefore, additional mechanisms of OPTN dysfunction must be associated with CNS axonopathy. Indeed, we found that OPTN binds directly to microtubules through a C-terminus-dependent mechanism (**Fig. 4A-C**). Consistently, the AlphaFold-Multimer predicates confident interaction between OPTN C-terminus and tubulin but very low interaction after C-terminus removal (**Fig. 4D**), providing a structural basis for understanding OPTN’s role in microtubule-dependent mitochondrial transport. Moreover, OPTN interacts with the mitochondria axonal anterograde transport complex, TRAK1-KIF5B-mitochondria, through a mechanism independent of the C-terminus (**Fig. 3F-I**). These interactions increase the probability that the TRAK1-KIF5B-mirochondria complex binds to the microtubule surface, which will increase both the chance that mitochondria will be transported along the microtubule and the distance that they travel, and therefore facilitates ample delivery of axonal mitochondria. Based on the neuron-autonomous degeneration caused by OPTNΔC in mouse RGCs and motor neurons, we propose that the C-terminus of OPTN is crucial in maintaining axon integrity, likely through a previously unknown function of OPTN in delivering axonal mitochondria that depends on the C-terminus (**Fig. 7C**). We have shown that loss of this function (OPTNΔC) significantly decreases axonal mitochondrial transport in *in vitro* reconstitution assays, in cultured neurons, and in *in vivo* mouse ONs. It is highly insightful that not only that C-terminus loss causes OPTN loss-of-function, but also that trapping of TRAK1-KIF5B-mtochondria by OPTNΔC has the potential to further decrease the binding of mitochondria to microtubules and therefore jeopardize microtubule-based transport of axonal mitochondria. This notion is further supported by the significant rescue of axonal mitochondria transportation/delivery and neurodegeneration in mouse retina/ON *in vivo* by overexpressing TRAK1/KIF5B in OPTNΔC RGCs.

Kinesin motor proteins have been long implicated in anterograde axon transport including in mitochondrial transport, and in neurodegenerative diseases in humans and in animal models including in RGC degeneration ^77–80^, our data extend this theme considerably by identifying the KIF5B engagement in OPTNΔC mechanism of degeneration. Intriguingly, we confirmed that ON mitochondria transport is also defective in ocular hypertension SOHU glaucoma mice, consistent with a recent publication using a different ocular hypertension model^24^. And excitingly, we found that overexpression of OPTN alone or combined with TRAK1/KIF5B dramatically increases survival of RGC somata and axons in this mouse glaucoma model. Therefore, deficient axonal mitochondria transport may be the common pathogenic feature of NTG and glaucoma with IOP elevation, which enhances the notion that targeting axonal mitochondrial transport represents a promising and potent neural repair strategy for both forms of glaucoma.

Here we demonstrated that overexpression of OPTN/TRAK1/KIF5B promotes striking ON regeneration that is comparable to that PTEN deletion but occurs through an independent and synergistic pathway. Support for this premise, strategies that augment axonal mitochondria transport are not only neuroprotective but also increase axon regeneration, is seen in prior work including deletion of the axonal mitochondria anchor protein syntaphilin and overexpression of Miro1 ^64, 65, 74^, expression of mitochondria protein ARMCX1 ^73^, and inhibition of HDAC6 ^75^. These data are in contrast to prior work in which overexpression of KIF5A alone is not sufficient to promote neuroprotection or regeneration ^79^, suggesting that co-expression of critical functional adaptors is also required to confer such significant phenotypes. Because many CNS neurodegenerative diseases have deficits in axonal mitochondria delivery ^15–24^, it is important to test whether exogenous OPTN/TRAK1/KIF5B expression increases neuroprotection and regeneration in animal models of these diseases as a general neural repair strategy for CNS axonopathies. It would also be worthwhile to perform high throughput screening to identify potent genes or small molecules that can significantly enhance axonal mitochondria delivery^81^ and would be promising candidates for novel neural repair therapies.

Several transgenic mouse lines containing the glaucoma-associated OPTN^E50K^ mutation have been generated before. Because the glaucomatous neurodegeneration phenotypes of these mice are heterogenous and appear only at a very late stage (e.g. in 12-18 months old mice) ^10, 82, 83^, however, they offered limited insights to the mechanisms of OPTN-induced neurodegeneration or to test potential therapeutic strategies. There have been even fewer reports of ALS animal models with OPTN mutation except a report that OPTN deletion in oligodendrocytes and microglia but not in neurons sensitizes these glial cells to necroptosis and causes nonautonomous neurodegeneration ^84^. The RGC specific OPTN C-terminus truncation mouse presented here represents an NTG-like animal model in which pronounced glaucomatous neuron-autonomous degeneration occurs rapidly and reliably. This OPTNΔC induced NTG-like model will certainly broaden the toolset of glaucoma research, especially for neuroprotection/neural repair. This proof-of-concept model with an NTG/ALS causative gene also warrants testing the effect of OPTN truncation in other neuronal cell types. Our Vglut2^+^-neuron OPTNΔC mouse is a promising model for ALS because it shows motor neuron degeneration and ALS-like locomotor deficits. This model could be improved by using more restricted motor neuron-specific Cre lines to mimic motor neuron-autonomous ALS. The two models reported here suggest that the OPTN C-terminus is critical for the proper functioning of OPTN in various CNS neurons, and that NTG and ALS share common pathogenic mechanisms.

Taken together, the OPTNΔC models of neurodegeneration that we establish and characterize here should prove to be invaluable tools for studying *in vivo* axonal mitochondria transport, NTG/ALS axonal pathology, and experimental therapies for neuroprotection and regeneration, and therefore for translating relevant findings into novel and effective treatments for patients with glaucoma, ALS, and other neurodegenerative diseases. The novel function of OPTN (**Fig. 7C**) in stabilizing the mitochondrial transport complex on microtubule to facilitate axonal mitochondria transport and the striking neuroprotection and axon regeneration phenotypes caused by OPTN/TRAK1/KIF5B point to a new mechanism of pathogenesis caused by OPTN dysfunction and a potent and promising neural repair strategy.

## Materials and Methods

### Mice

C57BL/6J WT (000664), Optntm1.1Jda/J (029708) “OPTN floxed” (OPTN^f/f^), Slc17a6tm2(cre)Lowl/J “Vglut2-Cre” (016963), B6.Cg-Gt(ROSA)26Sortm1(CAG-EGFP)Brsy/J “Mito-Tag” (032290), B6.129S4-Ptentm1Hwu/J “PTEN floxed” (006440), and B6.129S1-Kif5btm1Njen/J “KIF5B floxed” (008637) mice were purchased from Jackson Laboratories (Bar Harbor, Maine). Thy1-lsl-YFP1 ^85^ is a gift from Dr. Joshua Sanes’s lab. All mice were housed in standard cages on a 12-hour light-dark cycle. All experimental procedures were performed in compliance with animal protocols approved by the IACUC at Stanford University School of Medicine.

### Constructs and AAV production

Plasmids pLV-mitoDsRed (# 44386), pmCherry C1 Omp25 (#157758), and meGFP-hTRAK1 (# 188664) were purchased from Addgene. The coding regions of hOPTN (Addgene, #23053) and hOPTNΔC (Addgene, #23053) were cloned into meGFP-hTRAK1 to generate meGFP-hOPTN and meGFP- hOPTNΔC. The coding region of KIF5B ^71^, TRAK1 (Addgene, #127621), his-hOPTN (Addgene, #23053), mOPTN (mouse tissue cDNA), pMitoTimer (Addgene, #52659), 4xmts-mScarlet-I (Addgene, #98818), TurboID (Addgene, #107169) and Cre ^33^ were cloned and packaged into our pAM-AAV-mSncg-WPRE backbone containing the RGC-specific full-length mSncg promoter ^33^. Cre was cloned into pAM-AAV-mSncg-EGFP-WPRE to generate AAV-mSncg-Cre-T2A-EGFP. The detailed procedure of the AAV production has been described previously ^33^. Briefly, AAV plasmids containing the target genes were co-transfected with pAAV2 (pACG2)-RC triple mutant (Y444, 500, 730F) and the pHelper plasmid (StrateGene) into HEK293T cells by the PolyJet (SignaGen Laboratories, SL100688) transfection reagent. After transfection for 72 hours, the cells were lysed and purified by two rounds of cesium chloride density gradient centrifugation. The AAV titers of target genes were determined by real-time PCR and diluted to 1.5 x 10^12^ vector genome (vg)/ml for intravitreal injection.

### Ophthalmological procedures and measurements

The detailed procedures for intravitreal injection, IOP measurement, histological studies of RGC and ON, *in vivo* OCT imaging, PERG and OKR have been published before ^33, 39, 42, 86^. Brief descriptions are presented below.

### Intravitreal injection

For intravitreal injection, mice were anesthetized by an intraperitoneal injection of Avertin (0.3 mg/g) and 0.5% proparacaine hydrochloride (Akorn, Somerset, New Jersey) was applied to the cornea to reduce its sensitivity during the procedure. A pulled and polished microcapillary tube was inserted into the peripheral retina just behind the ora serrata. Approximately 2 µl of the vitreous was removed to allow injection of 2 µl AAV, MitoTracker Orange CMTMRos (Thermo Fisher Scientific, 0.15mM), cholera toxin subunit B-Alexa 555 or 647 (CTB555 or CTB647, Invitrogen, 2µg/µl) into the vitreous chamber. The mice were housed for an additional 2 weeks after AAV injection to achieve stable target genes expression. For anterograde tracing of mitochondria, MitoTracker Orange was injected 3 hours or CTB was injected 3 days before tissue collection and processing for imaging.

### Immunohistochemistry of retinal wholemounts and cross sections; RGC counts

After transcardiac perfusion with 4% PFA in PBS, the eyes were dissected out, post-fixed with 4% PFA for 2 hours, at room temperature, and cryoprotected in 30% sucrose overnight. For cryo-section with Leica cryostat, the eyeballs were embedded in Tissue-Tek OCT (Sakura) on dry ice for subsequent cryo-section.

For immunostaining of the wholemounts, retinas were dissected out and washed extensively in PBS before blocking in staining buffer (10% normal goat serum and 2% Triton X-100 in PBS) for 2 hours before incubating with primary antibodies: guinea pig anti-RBPMS 1:4000 (ProSci, California); rat anti-HA 1:200 (Roche, California); rabbit anti-OPTN C-terminus 1:1000 (Cayman Chemical, No. 100000), rabbit anti-OPTN polyclonal 1:1000 (Cayman Chemical, No. 100002), or Streptavidin-Alexa Fluor 488 conjugate 1: 200 (Invitrogen, CA) overnight at 4°C. After washing 3 times for 30 minutes each with PBS, samples were incubated with secondary antibodies (1:400; Jackson ImmunoResearch, West Grove, Pennsylvania) for 1 hour at room temperature. Retinas were again washed 3 times for 30 minutes each with PBS before a cover slip was attached with Fluoromount-G (SouthernBiotech, Birmingham, Alabama). Images of immunostained wholemounts were acquired with a Keyence epifluorescence microscope (BZ-X800) or Zeiss confocal microscope (LSM 880) with 20x and 40x oil lens. For RGC counting, 8 circles drawn by Concentric Circle plugin of NIH Fiji/ImageJ were used to define the peripheral, middle, and inner areas of the retina. Multiple 250 × 250 μm counting frames were applied by Fiji/ImageJ and the number of surviving RGCs was counted by RGCode software ^87^. The percentage of RGC survival was calculated as the ratio of surviving RGC numbers in treated eyes compared to contralateral control (CL) eyes. The investigators who counted the cells were blinded to the treatment of the samples.

### Optic nerve (ON) semi-thin sections and quantification of surviving axons

Transverse semi-thin (1 µm) sections of ON were cut on an ultramicrotome (EM UC7, Leica, Wetzlar, Germany) from tissue collected 2 mm distal to the eye and stained with 1% para-phenylenediamine (PPD) in methanol: isopropanol (1:1) solution. ON sections were imaged and stitched through a 100x lens of a Keyence fluorescence microscope. Multiple 10 × 10 μm counting frames were applied automatically by AxonCounter plugin of Fiji/ImageJ to sample about 10% of each ON. The number of surviving axons in the sampled areas was manually identified and counted. The mean of the surviving axon number was calculated for each ON and compared to that in the contralateral control ON to yield a percentage of axon survival value. The investigators who counted the axons were masked to the treatment of the samples.

### SOHU glaucoma model and IOP measurement

The detailed procedure has been published before ^38–41^. Briefly, mice were anesthetized by an intraperitoneal injection of Avertin (0.3 mg/g) and received the SO (1000 mPa.s, Silikon, Alcon Laboratories, Fort Worth, Texas) injection at 9 weeks of age. Prior to injection, 0.5% proparacaine hydrochloride (Akorn, Somerset, New Jersey) was applied to the cornea to reduce its sensitivity during the procedure. A 32 G needle was tunneled through the layers of the cornea at the superotemporal side close to the limbus to reach the anterior chamber without injuring lens or iris. Following this entry, ∼2 µl silicone oil was injected slowly into the anterior chamber using a sterile glass micropipette, until the oil droplet expanded to cover most areas of the iris (diameter ∼1.8–2.2 mm). After the injection, veterinary antibiotic ointment (BNP ophthalmic ointment, Vetropolycin, Dechra, Overland Park, Kansas) was applied to the surface of the injected eye. The contralateral control eyes received mock injection with 2 µl normal PBS to the anterior chamber. Throughout the procedure, artificial tears (Systane Ultra Lubricant Eye Drops, Alcon Laboratories, Fort Worth, Texas) were applied to keep the cornea moist.

The IOP of both eyes was measured by the TonoLab tonometer (Colonial Medical Supply, Espoo, Finland) according to product instructions. Briefly, mice were anesthetized and 1% Tropicamide sterile ophthalmic solution (Akorn, Somerset, New Jersey) was applied three times at 3-minute intervals to fully dilate the pupils before taking measurements. The average of six measurements by the TonoLab was considered as one machine-generated reading and three machine-generated readings were obtained from each eye; the mean was calculated to determine the IOP. During this procedure, artificial tears were applied to keep the cornea moist.

### Optic nerve crush (ONC)

ONC was performed 2 weeks following AAV injection when mice were about 7–8 weeks of age. After anesthetization by intraperitoneal injection of Avertin (0.3 mg/g), the ON was exposed intraorbitally, while care was taken not to damage the underlying ophthalmic artery, and crushed with a jeweler’s forceps (Dumont #5; Fine Science Tools, Foster City, CA, USA) for 5 seconds approximately 0.5 mm behind the eyeball. Eye ointment containing neomycin (Akorn, Somerset, New Jersey) was applied to protect the cornea after surgery.

### CTB tracing in wholemount ON and imaging

Intravitreal injection of CTB was performed 48 h before perfusion of the animals with 4% PFA in PBS. The ONs were carefully dissected with fine forceps and scissors and cleared with a modified iDISCO method ^88^, washed with PBS for 4 × 30 min; then immersed in a series of 20%, 40%, 60%, 80%, and 100% methanol in PBS for 30 min at each concentration for dehydration; dichloromethane (DCM)/methanol (2:1) for 30 min; 100% DCM for 30 min for clearance and dibenzyl ether (DBE) for 10 min before mounting on slides. Tiled images of the wholemount ON were captured and stitched by a Zeiss LSM 880 confocal laser scanning microscope with 25x/1.0 Oil lens (Carl Zeiss Microscopy, Thornwood, NY, USA), with Z-stack Airy scan. The number of CTB labeled axons was quantified as described previously^43^. Briefly, the fibers were counted that crossed perpendicular lines distal to the crush site every 250 μm till no fibers were visible. The average density (axon number/ μm^2^) from three sampled stacks (Z = 60, 120 and 180μm) was utilized to estimate the total number of axons using the formula ∑ad = πr² * mean axon density, where r is the optic nerve’s radius. All CTB signals shown in maximal projection that was set from lowest intensity to the maximum intensity after background subtraction were counted as individual fibers.

### Brain-optic nerve clearance and light sheet microscopy imaging

The whole brain with ONs and eyeballs was carefully dissected and embedded in 0.5% agarose gel block. The embedded gel block was cleared with a modified iDISCO method ^88^: PBS for 4 hours; a series of 20%, 40%, 60%, 80%, and 100% methanol in 1xPBS for 1day at each concentration; dichloromethane (DCM)/methanol (2:1) for 1 day; 100% DCM for 1day and dibenzyl ether (DBE) for 1 day. The ventral side of the tissue gel block was faced up and fixed on a spike holder, then placed into the imaging chamber immersed in the DBE buffer. The Light Sheet Ultramicroscope II generated 6 bi-directional 3.89μm thin light sheets to illuminate the tissue gel block from both sides while imaging the excited plane with a 2x objective microscope perpendicular to the sample. Tissue was imaged with the diode 561nm and 639nm laser and sheet numerical aperture (NA) 0.149 through a 2μm step-size of the Z-stack. The multiple optical sliced images of the whole tissue were collected and reconstructed by Imaris software (Oxford Instruments).

### Spectral-domain optical coherence tomography (SD-OCT) *in vivo* imaging

Fundus OCT imaging was performed under OCT mode by switching to a 30° licensed lens (Heidelberg Engineering), as previously described ^38, 42^. Briefly, the mouse retina was scanned with the ring scan mode centered by the ON head at 100 frames average under high-resolution mode (each B-scan consisted of 1536 A scans). The average thickness of GCC (includes retinal nerve fiber layer, ganglion cell layer, and inner plexiform layer) around the ON head was measured with the Heidelberg software (Heidelberg Engineering, Franklin, MA). The mean of the GCC thickness in the treated retina was compared to that in the contralateral control (CL) retina to yield a percentage of GCC thickness value. The investigators who measured the thickness of GCC were blinded to the treatment of the samples.

### Pattern electroretinogram (PERG) recording

PERG recording of both eyes was performed with the Miami PERG system (Intelligent Hearing Systems, Miami, Florida). A feedback-controlled heating pad (TCAT-2LV, Physitemp Instruments Inc., Clifton, New Jersey) maintained animal core temperature at 37°C. A small lubricant eye drop (Systane) was applied before recording to prevent corneal opacities. The reference electrode was placed subcutaneously on the occiput between the two ears, the ground electrode was placed at the root of the tail and the active steel needle electrode was placed subcutaneously on the snout for the simultaneous acquisition of left and right eye responses. Two 14 cm x 14 cm LED-based stimulators were placed in front so that the center of each screen was 10 cm from each eye. The pattern remained at a contrast of 85% with a luminance of 800 cd/m^2^, and consisted of four cycles of black-gray elements, with a spatial frequency of 0.052 c/d. Upon stimulation, the independent PERG signals were recorded from the snout and simultaneously by asynchronous binocular acquisition. With each trace recording up to 1020 ms, two consecutive recordings of 100 and 300 traces were averaged to achieve one readout. The first positive peak in the waveform was designated as P1 and the second negative peak as N2. The mean amplitude of the P1-N2 amplitude in the injured eye was compared to that in the contralateral control eye to yield a percentage of amplitude change. The investigators who measured the amplitudes were blinded to the treatment of the samples.

### Optokinetic response (OKR) measurement of visual acuity

The spatial vision of both eyes was measured using the OptoMotry system (CerebralMechanics Inc., Lethbridge, Alberta, Canada) dependent on opto-kinetic response (OKR). In brief, mice were placed unrestrained on a platform in the center of four 17-inch LCD computer monitors (Dell, Phoenix, Arizona); their movement was captured by a video camera above the platform. A rotating cylinder with vertical sine wave grating was computed and projected to the four monitors by OptoMotry software (CerebralMechanics Inc., Lethbridge, Alberta, Canada). The sine wave grating provides a virtual-reality environment to measure the spatial acuity of left eye when rotated clockwise and right eye when rotated counterclockwise. When the mouse calmed down and stopped moving, the gray of the monitor immediately switched to a low spatial frequency (0.1 cycle/degree) for five seconds, in which the mouse was assessed by judging whether the head turned to track the grating. The mice were judged to be capable of tracking the grating. The spatial frequency was increased repeatedly until a maximum frequency was identified and recorded. The % of vision acuity was yielded by comparing the maximum frequency of the experimental eye to that of the contralateral eye. The investigator who judged the OKR was blinded to the treatment of the mice.

### Mouse motor neuron behavioral test

All the mouse behavioral tests were performed at Stanford Behavioral and Functional Neuroscience Laboratory following the Stanford Operation Procedures (SOPs) for mouse behavioral testing. Four and 12-week-old mice were used for all the behavioral experiments and the ratios of males to females were approximately 1:1 in OPTN^f/f^ and OPTN^f/f^::Vglut2-Cre groups. Prior to the test, mice in home cage are placed in testing room for at least 1 hour before testing to minimize effects of stress on behavior during testing. Researchers were blinded to mouse genotypes.

### Grip strength test

The grip strength test is designed to assess motor function and control of the fore and hind paws. Mice were allowed to grab the bar on the Chatillon DFIS-10 digital force gauge (Largo, Florida USA) while being gently pulled parallel away from the bar by the tail. The maximum force prior to release of the mouse’s forepaws and hind paws from the bar was recorded. At least 20 minutes were allowed between each trial and 3 trials were taken for each mouse. After each trial, the apparatus was cleaned with a 1% Virkon solution. Maximum force exerted was noted (in Newtons (N)) and divided by the body weight to get force/body weight (N/g).

### Open field test

Locomotor activity as part of the motor neuron function was evaluated using the Open Field Test. The Open Field Activity Arena (43 x 43x 30 cm, Model ENV-515, Med Associates Inc, St. Albans, Vermont, USA) contains three planes of 16 infrared photobeam sensors, within a sound-attenuating and ventilated chamber (74 x 60 x 60 cm, MED-017M-027, Med Associates Inc, St. Albans, Vermont, USA). For testing, the mouse was placed in a corner of the arena and allowed to move freely for 30 min monitored by an automated tracking system (Activity Monitor Version 7, SOF-812, Med Associates Inc, St. Albans, Vermont, USA). The trial began immediately and ended when the defined duration had elapsed. Arena was cleaned with 1% Virkon between trials. Total distance traveled in the test was recorded.

### Rotarod test

The accelerating rotarod test was used to evaluate motor alterations in all the genotypes. Briefly, each mouse to be tested was pre-trained initially to stay on a no-accelerating rod (4rpm) elevated 16.5 cm above the testing floor for 1 min (Model ENV-575M, Med Associates Inc, St. Albans, Vermont, USA). For test session, all the mice were examined in an accelerating rod (4 to 40 rpm, with a cut off of 300 sec). The test session began when acceleration was started and ended when the animal fell off the rod. The mice were tested for three trials with 15-20 min inter-trial-intervals (ITIs). A 4^th^ trial was tested on the mice that held onto the rod for 2 consecutive revolutions or fell within 5 sec of the start of a trial. The apparatus is cleaned with 1% Virkon solution between trials. The 3 highest scores of the latency to fall from the rod during the testing session were recorded, and the average of the latency to fall for 3 trials was used for the analysis.

### Immunohistochemistry of spinal cord sections and motor neuron counts

Spinal cords fixed and embedded in Tissue-Tek OCT were sectioned in 30-μm-thicksections with cryo station CM3050. Sections were washed in PBS and blocked with 5% normal horse serum (VectorLabs). Primary antibodies Rabbit anti-NeuN (1:300, Proteintech), Goat anti-ChAT (1:200, Millpore) diluted in PBS supplemented with 2% horse serum and 0.1% triton X-100 (Sigma-Aldrich) were applied onto the sections followed by overnight incubation at 4°C in a humidified chamber. The slides were then washed in PBS and incubated with the appropriate secondary antibody diluted in PBS containing 2% horse serum and 0.1% triton X-100 (Sigma-Aldrich) with Donkey anti-rabbit Alexa488 (1:200 Jackson ImmunoResearch, Newmarket, UK) and Donkey anti-goat Cy3 (1:200 Jackson ImmunoResearch, Newmarket, UK) together for 2 h at room temperature, washed 3 times with PBS and mounted with DAPI Fluoromount-G (Southern Biotech, Birmingham, Alabama). Images of lumbar spinal cord were acquired with an Olympus Confocal Laser Scanning Microscope with FV3000 with 10x and 20x lens. For motor neuron identification and counting, spinal motor neurons were counted between lumbar segments 1–3, to minimize spatial effects on motor neuron counting. Briefly, images of spinal cord ventral horn were analyzed with Imaris software (Oxford Instruments) and NeuN positive neurons larger than 500um^2^ were selected and counted manual for both sides. ChAT positive neurons were manually identified and counted with Fiji/ImageJ. The investigators who counted the cells were blinded to the treatment of the samples.

### Mitochondria quantification in ON longitudinal sections and retinal wholemounts

ON longitudinal sections (8 µm) from globe to chiasm and retinal wholemounts labeled with 4xMTS- Scarlet, MitoTracker Orange, MitoTimer, Scarlet or CTB555 were imaged on a Zeiss confocal microscope (LSM 880) with a 40x lens for retina and 20x lens for ON. 4-8 images along the mid-to peripheral retina or 6-8 images from at least 3 separate sections from proximal to distal along each ON were taken with Z-stack images (7-8µm). Images were saved as maximum intensity projections with Fiji/Image for quantification of total fluorescence intensity. Confocal images from a single channel (MitoTracker Orange, 4xMTS-Scarlet, Scarlet, or CTB555) were imported into Fiji/ImageJ and the Corrected Total Fluorescence (CTF) was quantified: CTF = Integrated Density of the entire image – (Area of image x Mean fluorescence of background readings) (Luke Hammond, QBI, The University of Queensland, Australia: https://theolb.readthedocs.io/en/latest/imaging/measuring-cell-fluorescence-using-imagej.html). The mean CTF value was calculated for each retina wholemount and ON section and compared to that in the contralateral control to yield a percentage of CTF value. For MitoTimer Red:Green ratio, the CTF mean values for each retinal wholemount and ON section were quantified for both channels (EGFP and DsReD). For mitochondrial morphology quantification, high resolution ON longitudinal images labeled with 4MTS-Scarlet were analyzed by the ImageJ plugin MitoMap (http://www.gurdon.cam.ac.uk/stafflinks/downloadspublic/imaging-plugins) ^89^. MitoMap automates the process of defining Scarlet-labeled mitochondria in a selected region of interest and calculates their volume, surface area, and shape descriptors including sphericity (ratio of the surface area of a sphere with the same volume as the object to the surface area of the object), distribution isotropy (sum of ratios of the second moments in each combination of orientations) and compactness (variance of radial distance/volume) of mitochondria for both groups.

### ON wholemount clearance and quantification for mitochondrial density

The ONs labeled with MitoTracker Orange were trimmed and cleared by a modified iDISCO method ^88^: samples were incubated with PBS for 30mins, and then a series of 20%, 40%, 60%, 80%, and 100% methanol in 1xPBS for 30mins at each concentration before dichloromethane (DCM)/methanol (2:1) for 30mins; 100% DCM for 30mins, and dibenzyl ether (DBE) for 30mins. The cleared ONs were incubated with DBE on slides between two 22 x 22 mm cover slips, covered with a 22 x 30 mm cover slip, and sealed with clear nail polish. The mounted ON wholemounts were imaged with a 10x objective lens using the tile scan for the whole length of the ONs, and 40x oil immersion objective lens with three Z stack images (100µm x 100µm, n = 50, Z interval = 0.5µm) per ON at three locations (proximal, ∼1.5mm; distal, 3.0mm; distal, 4.5mm) distance to the eyeball. Each Z-stack image series was imported to Imaris Software (Oxford Instruments) and reconstructed to 3-dimensional structure for mitochondrial quantification with the spots tool in Imaris using the same threshold parameters (diameter ∼ 1.5 µm). Manual adjustment was made to cover all the mitochondria in the field. The total number of mitochondria in Z stack images of the treated eyes was compared to that in the contralateral control eyes to yield a percentage of Mito Density.

### Transmission Electron Microscope (TEM) imaging and quantification of mitochondria in ON ultrathin cross-sections

Ultrathin cross-sections of the ON 2 mm distal to the eye (globe) were collected and stained with uranyl acetate for 30 minutes, washed in PBS and then stained with lead citrate for 7 minutes. Sections were again washed and dried before observing under TEM. The cross-sections of the entire ON were examined and imaged randomly without overlap at 10,000× on a JEOL JEM-1400 TEM microscope (JEOL USA, Inc., Peabody, MA). For each ON, 10-16 images per ON were taken to cover the area of ON. The total number of mitochondria was counted and divided by the number of total axons in the same image to get the #mitochondria/axon per image. Aspect ratio was quantified by measuring the full width of a mitochondrion in Fiji/ImageJ and dividing that by the full height of that mitochondrion, on all mitochondria per ON images and combined per treatment group. Mitochondria area was quantified by using an ellipses area equation: Area = π*a*b where a is the height radius and b is the length radius of the mitochondria.

### *Ex vivo* time-lapse imaging of ON mitochondria

To image axonal transport of mitochondria in optic nerve, 2 µL of MitoTracker Orange (0.15mM) was intravitreally injected into the vitreous chamber 3h before ex vivo time-lapse imaging. The optic nerve were quickly harvested at 2 weeks post injection of AAV-Cre or Capsid into OPTN^f/f^ mice, or at 3 weeks post induction in the SOHU model of AAV-mOPTN or mOPTN+TRAK1+KIF5B mice and maintained in Hibernate E low-fluorescence medium (BrainBits) at 37°C on a heated stage and further transferred to 35mm glass-bottom dishes (MatTek) pre-coated with poly-L-lysine (0.5 mg/ml in ultrapure water) with coating medium (methyl cellulose 8mg/ml in Hibernate E medium). Time-lapse images were captured through a 40x oil immersion objective lens at 1 frame per 2 s for 5 min using a Zeiss LSM880 confocal microscope equipped with an incubation chamber. Mitochondrial events were traced, and kymograph analyses were performed using Kymolyzer plugin of Fiji/ImageJ software ^90^. Briefly, mitochondria with average instantaneous velocity higher than 0.05µm/s were categorized as motile. Mitochondria with average instantaneous velocity lower than 0.05µm/s were considered as zero ^91^. The following parameters were determined using Kymolyzer plugin: 1) percentage of mobile time of each mitochondrion; 2) percentage of stationary time of each mitochondrion; 3) average speed of each mitochondrion is in motion; 4) move length of each mitochondrion is in motion; and 5) percentage of mitochondria in motion.

Mitochondrial density was determined by manually counting the total number of mitochondria from each 100-µm-long distalmost axonal segment using Fiji/ImageJ software (NIH).

### Hippocampal neuron culture and time-lapse imaging acquisition and quantification

Hippocampal cells were dissociated from day 15 OPTN^f/f^ or wild-type (WT) mouse embryos, cultured in 35mm glass-bottom dishes (MatTek) pre-coated with poly-L-lysine (0.5 mg/ml in ultrapure water) with Neurobasal Plus culture medium with 2% B27 (Gibco) for 5 days, and then transfected with Lipofectamine 3000 (Invitrogen) with EGFP-OPTN, pLV-MitoDsRed or pmCherry C1-OMP25 for WT hippocampal neurons or AAV-mSncg-Cre-T2A-EGFP, AAV-mSncg-EGFP ^33^, or pLV-MitoDsRed for OPTN^f/f^ hippocampal neurons. Twenty-four hours post transfection, time-lapse confocal images were acquired with a 40x oil immersion objective lens at 1 frame per 2 s for 5 min using a Zeiss LSM880 confocal microscope equipped with an incubation chamber maintained at 37°C with 5% CO2. Mitochondrial events were traced and kymograph analyses were performed using Kymolyzer plugin of Fiji/ImageJ as described above.

### 3D-structured illumination super-resolution microscope (SIM) imaging

E15 DIV hippocampus neurons described above were transfected with EGFP-OPTN or EGFP-OPTNΔC 24 hrs before staining with SPY555-tubulin (Cytoskeleton. Inc) for SIM imaging. The images were acquired using a DeltaVision OMX Blaze imaging system (GE Healthcare) equipped with U-Planapo 100X SIM lens, 3 channels emCCD cameras, Piezo controlled Fast Z-axis system (100 um range) and 488, 568 nm MONET lasers for excitation. The SIM set-up uses BLAZE SI patterns and acquires images per 3 µm z-stack thickness with 0.125 µm section spacing (3 illumination angles times 5 phase pixel size 0.0807 µm x 0.0807 µm x 0.125 µm) per color channel. Super-resolution 3D images are then obtained via image processing using the reconstruction software. Image deconvolution and 3DSIM reconstructions were completed using the manufacturer-supplied softWoRx program (GE Healthcare). Image registration (color channel alignment) was also performed in the same program using experimentally-measured calibration values compensating for minor lateral and axial shifts, rotation, and magnification differences between cameras. Image analysis and processing after the deconvolution and alignment was done using Imaris software, including the conversion from DV to TIFF image files (preserving bit-depth and metadata) and visualization using orthogonal views.

### Proximity labeling with OPTN-TurboID in HEK293 cells

PolyJet (SignaGen Lab) was incubated with serum-free Dulbecco’s Modified Eagle Medium (DMEM, GIBCO) and used to transfect HEK293 cells with a mixture of OPTN-TurboID plasmid. Twenty-four hours after transfection, we added exogenous 500 μM of biotin (Thermo Scientific) to the culture medium, which was from a 100 mM biotin stock diluted in dimethyl sulfoxide (DMSO) and incubated for 24 hours at 37 ℃. To stop the labeling, cells were transferred on ice and washed five times with ice-cold PBS for 10 seconds each wash. The OPTN-TurboID expressing HEK293 cells were treated with or without biotin, and naïve cells were lysed with RIPA lysis and extraction buffer supplemented with complete protease inhibitor (Thermo Scientific), studied for western blotting with rabbit anti-OPTN polyclonal 1:1000 (Cayman Chemical, No. 100002) or Streptavidin-HRP 1:20,000 (Invitrogen) and examined by chemiluminescence using ECL (Thermo Fisher Scientific, Massachusetts).

### *In vivo* proximity labeling with OPTN-TurboID in RGCs

The 5-week-old C57BL/6 mice were injected with AAV2-mSncg-OPTN-TurboID or TuboID intravitreally. At 4 weeks post AAV injection, the mice were intravitreally injected with 70 mM of biotin 24 hours before sacrifice and retina collection. The fresh retinas were washed twice with cold PBS before lysis. For each condition, 16 retinas were pooled and resuspended in lysis buffer (50 mM Tris pH 7.4, 500 mM NaCl, 0.4% SDS, 5 mM EDTA, 1 mM DTT, 1x complete protease inhibitor), and then passed 10-20 times through a 19-G needle before four cycles of sonication at 30% intensity, 30 seconds per cycle in cold water bath. Triton X-100 was added to the recovered sonicated lysate to reach a final concentration of 2% before adding 50 mM Tris to adjust the NaCl concentration to 150 mM before binding to streptavidin-coupled beads. The adjusted lysates were centrifuged at 16,000 xg, at 4°C for 10 min. The concentration of each sample was measured with a BCA colorimetric assay (ThermoFisher) and 3 mg protein lysate used for streptavidin pulldown with 200 μL of streptavidin-coupled magnetic beads (ThermoFisher, 65002). The beads were washed by gently mixing with: 50 mM Tris pH 7–4, 150 mM NaCl, 0.05% Triton X-100, 1 mM DTT. Then each set of beads was resuspended with equal amounts of the corresponding retina lysates and incubated overnight at 4 °C on a rotating wheel. On the next day, the beads were serially washed twice each for 8 min on a rotation wheel with: 2% SDS in water; 50 mM HEPES pH 7.4, 1 mM EDTA, 500 mM NaCl, 1% Triton X-100, and 0.1% Na-deoxycholate; 10 mM Tris pH 8, 250 mM LiCl, 1 mM EDTA, 0.5% NP-40, and 0.5% Na-deoxycholate; 50 mM Tris pH 7.4, 50 mM NaCl, 0.1% NP-40. Then the beads were washed four times for 5 min on a rotation wheel with 1x PBS. 40 μL of elution buffer (10 mM Tris pH 7.4, 2% SDS, 5% β-mercaptoethanol, and 2 mM Biotin) was added to the beads and incubated at 98 °C for 15 min, then the beads were immediately removed on a magnetic rack. The eluted samples were transferred and submitted to Stanford University Mass Spectrometry Core Facility for protein detection by LC-MS/MS, at Vincent Coates Foundation Mass Spectrometry Laboratory. Heatmap was generated using the R package ComplexHeatmap to visualize the enriched OPTN-interacting proteins in RGCs identified by *in vivo* TurboID.

### Co-immunoprecipitation (Co-IP)

Total HEK293 cell extracts were prepared according to the manufacture’s protocol (Pierce HA-Tag Magnetic IP/Co-IP kit and Pierce Protein A/G Magnetic Beads, Thermo Fisher Scientific, MA). Twenty-four hours after transfection, HEK293 cells were harvested, washed in ice-cold PBS, and lysed in 300 μl lysis/wash buffer (0.025M Tris, 0.15M NaCl, 0.001M EDTA, 1% NP40, 5% glycerol) containing 1x Halt Protease and Phosphatase Inhibitor Cocktail (Thermo Fisher Scientific, Massachusetts). For Anti-HA IP, extracted protein was immunoprecipitated using 25μL prewashed HA Ab-Tag Magnetic beads (Thermo Fisher Scientific, Massachusetts) with gentle rotation at 4°C for 30min. For Anti-His IP, extracted protein was combined with 3 µg His antibody overnight at 4°C with mixing, then mixed with prewashed Protein A/G Magnetic Beads (Thermo Fisher Scientific, Massachusetts) with gentle rotation for 3 hours at 4 °C. Immunoprecipitants were washed four times in lysis/wash buffer, and bound proteins were dissociated in 50 μL of 1x loading dye (0.3M Tris·HCl, 5% SDS, 50% glycerol, lane marker tracking dye). Eluted proteins were separated on SDS -12% polyacrylamide gel and transferred onto Supported Nitrocellulose Membrane (Biorad) or PVDF Membrane (Millipore). To prevent nonspecific binding, membranes were incubated in blocking buffer (5% skimmed dried milk, 20 mM Tris, 150 mM NaCl, 0.1% Tween-20) with agitation for 1 hour at room temperature, followed by immediate incubation with Rabbit anti-OPTN (1:1000, Proteintech), Rabbit anti-KIF5B (1:1000, Cell Signaling Technology), Rabbit anti-TRAK1( 1:1000, Invitrogen), Chick anti-GFP (1:1000, Aves Lab), or Rabbit anti-GAPDH (1:1000, Cell Signaling Technology) diluted in 5% BSA overnight. Membranes were then washed three times in washing buffer (20 mM Tris, 150 mM NaCl, 0.1% Tween-20), incubated for 1.5 hours at room temperature with goat anti-rabbit HRP-conjugated antibody or donkey anti-chick HRP-conjugated antibody (Thermo Fisher Scientific, Massachusetts). Protein expression was detected by chemiluminescence using ECL (Thermo Fisher Scientific, Massachusetts).

### Mitochondrial immunoprecipitation

Isolation of mitochondria was performed as previously described ^92, 93^. Briefly, transfected HET293T cells or MitoTag retinas were harvested and washed in ice-cold PBS before homogenization in 1x KPBS (136 mM KCl, 10 mM KH2PO4, pH 7.25). Five microliters of this homogenate were taken as input and extracted in Triton lysis buffer. The remaining homogenate was spun down at 1,000 × g for 2 min at 4 °C. The supernatant was subjected to immunoprecipitation with prewashed anti-HA beads (Thermo Scientific) for 30 mins, followed by three rounds of washing in 1x KPBS. In the final wash, proteins were extracted by mixing the beads with 50 μL of Triton lysis buffer and incubated on ice for 10min. The lysate was centrifuged at 17,000 x g for 10 min at 4 °C. The supernatant was mixed with loading dye and separated on SDS -12% polyacrylamide gel and transferred onto PVDF Membrane (Millipore) before incubation with rabbit anti-OPTN (1:1000, Proteintech), rabbit anti-Miro1 (1:1000, Novus biologicals), or rabbit anti-HA (1:1000, Cell Signaling Technology) diluted in 5% BSA overnight. Protein expression was detected by chemiluminescence using ECL after the membrane was incubated with goat anti-rabbit HRP-conjugated antibody.

### Production of recombinant proteins

N-terminally histidine- and mNeonGreen-tagged full length and truncated human optineurin (mNG- OPTN and mNG-OPTNΔC) was expressed in SF9 insect cells using the open source FlexiBAC baculovirus vector system ^94^. Cells expressing the protein were lysed in 50 mM Na-phosphate buffer pH 7.5, containing 300 mM KCl, 1 mM MgCl2, 5% glycerol, 0.1% Tween-20, 10 mM β-mercaptoethanol (BME), 0.1 mM ATP, 30 mM imidazole, Protease Inhibitor Cocktail (complete, EDTA free, Roche), 25 units/ml benzonase, and centrifuged at 30,000 G for 1 hour at 4 °C to remove the pellet. The supernatant was collected and loaded onto a Ni-NTA column. The column was washed with 50 mM Na-phosphate buffer pH 7.5, containing 300 mM KCl, 1 mM MgCl2, 5% glycerol, 0.1% Tween-20, 10 mM BME, 0.1 mM ATP, and 60 mM imidazole. Protein was eluted using 2 ml of 50 mM Na-phosphate buffer pH 7.5, 300 mM KCl, 1 mM MgCl2, 5% glycerol, 0.1% tween-20, 10 mM BME, 0.1 mM ATP, and 375 mM imidazole. The protein was dialyzed into 50 mM Na-Phosphate buffer of pH 7.5, containing 300mM KCL, 5% Glycerol, 1 mM MgCl2, 2.5 mM DTT, 0.1 mM ATP, and 0.05 % Tween 20 and concentrated in Amicon centrifugal filter 50K (Millipore) to a final volume of 500 uL. Protein was aliquoted to smaller aliquots, flash frozen and kept at -80 °C (for SDS gel see **Fig. S4A**). Expression and purification of TRAK1-mCherry and KIF5B was carried out as described previously ^71^. For the cell lysate experiments, OPTN or OPTNΔC overexpressing SF9 cells were lysed in 50 mM Na-phosphate buffer pH 7.5, containing 300 mM KCl, 1 mM MgCl2, 5% glycerol, 0.1% Tween-20, 10 mM β-mercaptoethanol (BME), 0.1 mM ATP, Protease Inhibitor Cocktail (complete, EDTA free, Roche), and centrifuged at 30,000 G for 1 hour at 4 °C to remove the pellet. The supernatant was collected and flash frozen to smaller aliquots.

### *In vitro* reconstitution motility assays

Biotinylated microtubules were polymerized from 4 mg/ml tubulin in BRB80 (80 mM PIPES, 1 mM EGTA, 1 mM MgCl2, pH 6.9) supplemented with 1 mM MgCl2 and 1 mM GTP (Jena Bioscience, Jena, Germany) for 30 minutes at 37 °C prior to centrifugation at 18.000 x g for 30 min in a Microfuge 18 Centrifuge (Beckman Coulter) and resuspension of the microtubule pellet in BRB80 supplemented with 10 µM Taxol (Sigma Aldrich, T7191) (BRB80T).

Flow cells were prepared by attaching two cleaned and salinized by DDS (0.05% dichloro-dimethyl silane in trichloroethylene) or HMDS ((Bis(trimethylsilyl)amine)) glass coverslips (22 x 22 mm2 and 18 x 18 mm2; Corning, Inc., Corning, NY, USA) together using heated strips of parafilm M (Pechiney Plastic Packaging, Chicago, IL, USA). The flow cells were incubated with 20 µg/ml anti-biotin antibody (Sigma Aldrich, B3640) in PBS for 10 min and passivated by 1% Pluronic F-127 (Sigma Aldrich, P2443) in PBS for at least 1 hour. The flow cells were then washed with BRB80T, microtubules were introduced into the channel and immobilized on the antibodies, and unbound microtubules were removed by a flush of BRB80T. Immediately prior to the experiment, the solution was exchanged by motility buffer (BRB80 containing 10 µM Taxol, 10 mM dithiothreitol, 20 mM d-glucose, 0.1% Tween-20, 0.5 mg/ml casein, 1 mM Mg-ATP, 0.22 mg/ml glucose oxidase, and 20 µg/ml catalase).

To test if OPTN interacts with microtubules, mNG-OPTN or mNG-OPTNΔC diluted in motility buffer to the final concentration was flushed in the channel with microtubules. To test the OPTN-TRAK1 interaction or KIF5B and OPTN interaction, respective components were incubated on ice for 10 minutes before the mixture was diluted to the final concentration with motility buffer and flushed into the channel. Protein concentrations used in the assays are denoted in the respective figure legends.

For experiments quantifying motility parameters (**Fig. 4E,F and Fig. S5A,B**), mixture of i) 11 nM KIF5B, 34.5 nM mCherry-TRAK1 or ii) 11 nM KIF5B, 34.5 nM mCherry-TRAK1 and 1 µM mNG- OPTN or 1 µM mNG-OPTNΔC, were incubated on ice for 10 min, diluted in motility buffer supplemented with 100 mM KCl to the final concentration and flushed into the channel.

To test the interaction of OPTN/OPTNΔC lysates with the microtubules, lysates of mNG-OPTN or mNG-OPTNΔC-overexpressing SF9 cells diluted in motility buffer (BRB40 containing 10 µM Taxol, 10 mM dithiothreitol, 20 mM d-glucose, 0.1% Tween-20, 0.5 mg/ml casein, 1 mM Mg-ATP, 0.22 mg/ml glucose oxidase, and 20 µg/ml catalase) were added to the microtubules.

For experiments quantifying motility parameters in cell lysates (**Fig. S5C-F**), mixtures of i) 16 nM purified KIF5B and the mCherry-TRAK1 lysate or ii) 16 nM purified KIF5B combined with the mCherry- TRAK1 lysate and the mNG-OPTN/mNG-OPTNΔC lysate, were incubated on ice for 10 min, diluted in motility buffer supplemented with 100 mM KCl to the final concentration and flushed into the channel.

All imaging was performed using total internal reflection fluorescence (TIRF) microscopy on an inverted NIKON microscope equipped with Apo TIRF 100x Oil, NA 1.49, WD 0.12 mm or Apo TIRF 60x Oil, NA 1.49,WD 0.12 mm objective and PRIME BSI (Teledyne Photometrics) camera or CMOS camera (sCMOS ORCA 4.0 V2, Hamamatsu Photonics). The imaging setup was controlled by NIS Elements software (Nikon). Microtubules were imaged using Interference Reflection Microscopy (IRM). All movies were taken with an exposure time of 200 ms or 100 ms over the span of 1 minute with a frame rate of 2.5 or 5 frames per second with triggered acquisition for two channels and 5 or 10 for single channel imaging.

### Image Analysis - reconstitution experiments

All movies were analyzed manually using FIJI software ^95^. Kymographs were generated using the FIJI Kymograph Builder. Frequency of migration events were calculated as the number of detected molecules per microtubule length per second. The run time (cumulative time spent in a continuous directed motility excluding pauses) and run length (distance traversed by a molecule or a complex along a microtubule) were measured from kymographs using the FIJI measure tool. Velocity of continuous migration of a molecule or a complex was calculated as a ratio of run length and run time for each continuous run (excluding pauses). Run length and run time survival probabilities were estimated using Kaplan-Meier statistics in MATLAB - runs of molecules/complexes for which the beginning or end was not observed during the experiment were considered as censored events. To estimate the density of OPTN/OPTNΔC on microtubules for purified proteins or in a lysate, the background-subtracted mNG signal density on the microtubule was measured. T-tests were done using ttest function in MATLAB or from GraphPad Prism; log-rank test to measure p-values for Kaplan Meier estimation was done using https://www.statskingdom.com/kaplan-meier.html.

### AlphaFold protein-protein interaction predictions

To further understand the molecular basis of OPTN’s interaction with microtubules and its implications for neurodegenerative diseases, we employed AlphaFold2 ^96^. This advanced AI-based program predicts protein structures by assessing distances and angles between amino acid pairs, thus providing insights into potential binding sites and interactions. The primary focus was on evaluating the likelihood and nature of the interaction between OPTN and tubulin, and comparing it with the truncated variant, OPTNΔC. Firstly, the protein sequences of OPTN, OPTNΔC, and Tubulin alpha-1A were input into AlphaFold2 to generate predicted structural models. We then analyzed these models to identify potential binding sites. We run these experiments using AlphaFold-Multimer function ^97^. The prediction confidence is visualized as Predicted Aligned Error (PAE) plots. PAE plots are essentially heatmaps that display the predicted error between all pairs of amino acid residues in a protein.

### Statistical analyses

GraphPad Prism 9 was used to generate graphs and for statistical analyses. Data are presented as means ± s.e.m. Paired Student’s t-test was used for comparison of the two eyes of the same animals, unpaired t-test was used for two groups of animals and behavioral data analysis, and One-way ANOVA with post hoc test was used for multiple comparisons.

## Acknowledgements

We thank Dr. Ryan Leib, Kratika Singhal and Rowan Matney at Vincent Coates Foundation Mass Spectrometry Laboratory, Stanford University Mass Spectrometry for their great technical support with LC/MS analysis, Stanford Wu Tsai Neuroscience Microscopy Service for Light Sheet Ultramicroscope II imaging, Stanford Cell Science Imaging Facility for OMX BLAZE 3D-structured illumination microscope (SIM) imaging, Dr. Xinnan Wang for mitochondria motility assay in cultured neurons, Roopa Dalal for semi-thin sections, and Katerina Konecna and Tereza Smidova for purification of optineurin. We thank Drs. Xiaojing Gao, Alan Tessler, Wenjun Yan, and Hu lab members for critical discussion and reading the manuscript.

## Author contributions

Y.H., H.C.W., D.L., Y.X., F.B., M.P., and Z.L. designed the experiments. H.C.W. established the NTG- like animal model; D.L., I.Y., L.Liu, F.B., and L,Li established the ALS-like animal model; X.F., H.Y., P.L., M.Y., and H.H. participated in the collection of in vivo data; Y.X. and H.H. performed in vivo RGC TurboID assays; M.P., M.B., and Z.L. performed the in vitro reconstitution motility assays and data analysis; D.L. and F.B. led the biochemical and imaging characterization of OPTN-TRAK1-KIF5B- mitochondria interaction in cultured neurons and ex vivo ONs; C.C. did AlphaFold analysis; L.L. produced AAVs; S.H.S, X.D., D.W., A.L., Y.S. and J.L.G. provided reagents and equipment and participated in discussions. Y.H., D.L., F.B., M.P., and Z.L. prepared the manuscript with support from all the authors.

## Data and materials availability

All data are available in the main text or the supplementary materials.

## Supplementary Materials

Figs. S1 to S7 Movies S1 to S7

## References

1. Coleman, M.P. & Perry, V.H. Axon pathology in neurological disease: a neglected therapeutic target. Trends Neurosci 25, 532–537 (2002).

2. Raff, M.C., Whitmore, A.V. & Finn, J.T. Axonal self-destruction and neurodegeneration. Science 296, 868–871 (2002).

3. Fischer, L.R., et al. Amyotrophic lateral sclerosis is a distal axonopathy: evidence in mice and man. Exp Neurol 185, 232–240 (2004).

4. Nickells, R.W., Howell, G.R., Soto, I. & John, S.W. Under pressure: cellular and molecular responses during glaucoma, a common neurodegeneration with axonopathy. Annu Rev Neurosci 35, 153–179 (2012).

5. Anderson, D.R., Drance, S.M., Schulzer, M. & Collaborative Normal-Tension Glaucoma Study, G. Natural history of normal-tension glaucoma. Ophthalmology 108, 247–253 (2001).

6. Maruyama, H., et al. Mutations of optineurin in amyotrophic lateral sclerosis. Nature 465, 223–226 (2010).

7. Rezaie, T., et al. Adult-onset primary open-angle glaucoma caused by mutations in optineurin. Science 295, 1077–1079 (2002).

8. Wild, P., et al. Phosphorylation of the autophagy receptor optineurin restricts Salmonella growth. Science 333, 228–233 (2011).

9. Ryan, T.A. & Tumbarello, D.A. Optineurin: A Coordinator of Membrane-Associated Cargo Trafficking and Autophagy. Front Immunol 9, 1024 (2018).

10. Minegishi, Y., Nakayama, M., Iejima, D., Kawase, K. & Iwata, T. Significance of optineurin mutations in glaucoma and other diseases. Prog Retin Eye Res 55, 149–181 (2016).

11. Qiu, Y., et al. Emerging views of OPTN (optineurin) function in the autophagic process associated with disease. Autophagy 18, 73–85 (2022).

12. Chamberlain, K.A. & Sheng, Z.H. Mechanisms for the maintenance and regulation of axonal energy supply. Journal of neuroscience research 97, 897–913 (2019).

13. Cheng, X.T., Huang, N. & Sheng, Z.H. Programming axonal mitochondrial maintenance and bioenergetics in neurodegeneration and regeneration. Neuron 110, 1899–1923 (2022).

14. Misgeld, T. & Schwarz, T.L. Mitostasis in Neurons: Maintaining Mitochondria in an Extended Cellular Architecture. Neuron 96, 651–666 (2017).

15. Calkins, M.J., Manczak, M., Mao, P., Shirendeb, U. & Reddy, P.H. Impaired mitochondrial biogenesis, defective axonal transport of mitochondria, abnormal mitochondrial dynamics and synaptic degeneration in a mouse model of Alzheimer’s disease. Hum Mol Genet 20, 4515–4529 (2011).

16. Vicario-Orri, E., Opazo, C.M. & Munoz, F.J. The pathophysiology of axonal transport in Alzheimer’s disease. J Alzheimers Dis 43, 1097–1113 (2015).

17. Kanaan, N.M., et al. Axonal degeneration in Alzheimer’s disease: when signaling abnormalities meet the axonal transport system. Exp Neurol 246, 44–53 (2013).

18. Chang, D.T., Rintoul, G.L., Pandipati, S. & Reynolds, I.J. Mutant huntingtin aggregates impair mitochondrial movement and trafficking in cortical neurons. Neurobiology of disease 22, 388–400 (2006).

19. Trushina, E., et al. Mutant huntingtin impairs axonal trafficking in mammalian neurons in vivo and in vitro. Mol Cell Biol 24, 8195–8209 (2004).

20. Bilsland, L.G., et al. Deficits in axonal transport precede ALS symptoms in vivo. Proc Natl Acad Sci U S A 107, 20523–20528 (2010).

21. Magrane, J., Cortez, C., Gan, W.B. & Manfredi, G. Abnormal mitochondrial transport and morphology are common pathological denominators in SOD1 and TDP43 ALS mouse models. Hum Mol Genet 23, 1413–1424 (2014).

22. Baldwin, K.R., Godena, V.K., Hewitt, V.L. & Whitworth, A.J. Axonal transport defects are a common phenotype in Drosophila models of ALS. Hum Mol Genet 25, 2378–2392 (2016).

23. Ito, Y.A. & Di Polo, A. Mitochondrial dynamics, transport, and quality control: A bottleneck for retinal ganglion cell viability in optic neuropathies. Mitochondrion 36, 186–192 (2017).

24. Quintero, H., et al. Restoration of mitochondria axonal transport by adaptor Disc1 supplementation prevents neurodegeneration and rescues visual function. Cell reports 40, 111324 (2022).

25. Takihara, Y., et al. In vivo imaging of axonal transport of mitochondria in the diseased and aged mammalian CNS. Proc Natl Acad Sci U S A 112, 10515–10520 (2015).

26. Crish, S.D., Sappington, R.M., Inman, D.M., Horner, P.J. & Calkins, D.J. Distal axonopathy with structural persistence in glaucomatous neurodegeneration. Proc Natl Acad Sci U S A 107, 5196–5201 (2010).

27. Kimball, E.C., et al. The effects of age on mitochondria, axonal transport, and axonal degeneration after chronic IOP elevation using a murine ocular explant model. Experimental eye research 172, 78–85 (2018).

28. Vanhauwaert, R., Bharat, V. & Wang, X. Surveillance and transportation of mitochondria in neurons. Curr Opin Neurobiol 57, 87–93 (2019).

29. Zahavi, E.E. & Hoogenraad, C.C. Multiple layers of spatial regulation coordinate axonal cargo transport. Curr Opin Neurobiol 69, 241–246 (2021).

30. Rezaie, T. & Sarfarazi, M. Molecular cloning, genomic structure, and protein characterization of mouse optineurin. Genomics 85, 131–138 (2005).

31. Monavarfeshani, A., et al. Transcriptomic analysis of the ocular posterior segment completes a cell atlas of the human eye. Proc Natl Acad Sci U S A 120, e2306153120 (2023).

32. Munitic, I., et al. Optineurin insufficiency impairs IRF3 but not NF-kappaB activation in immune cells. J Immunol 191, 6231–6240 (2013).

33. Wang, Q., et al. Mouse gamma-Synuclein Promoter-Mediated Gene Expression and Editing in Mammalian Retinal Ganglion Cells. J Neurosci 40, 3896–3914 (2020).

34. Weil, R., Laplantine, E., Curic, S. & Genin, P. Role of Optineurin in the Mitochondrial Dysfunction: Potential Implications in Neurodegenerative Diseases and Cancer. Front Immunol 9, 1243 (2018).

35. Weishaupt, J.H., et al. A novel optineurin truncating mutation and three glaucoma-associated missense variants in patients with familial amyotrophic lateral sclerosis in Germany. Neurobiology of aging 34, 1516 e1519–1515 (2013).

36. Ozoguz, A., et al. The distinct genetic pattern of ALS in Turkey and novel mutations. Neurobiology of aging 36, 1764 e1769–1764 e1718 (2015).

37. Goldstein, O., et al. OPTN 691_692insAG is a founder mutation causing recessive ALS and increased risk in heterozygotes. Neurology 86, 446–453 (2016).

38. Zhang, J., et al. Silicone oil-induced ocular hypertension and glaucomatous neurodegeneration in mouse. eLife 8 (2019).

39. Fang, F., et al. Chronic mild and acute severe glaucomatous neurodegeneration derived from silicone oil-induced ocular hypertension. Scientific reports 11, 9052 (2021).

40. Zhang, J., et al. A Reversible Silicon Oil-Induced Ocular Hypertension Model in Mice. Journal of visualized experiments : JoVE 153 (2019).

41. Moshiri, A., et al. Silicone Oil-Induced Glaucomatous Neurodegeneration in Rhesus Macaques. Int J Mol Sci 23 (2022).

42. Li, L., et al. Longitudinal Morphological and Functional Assessment of RGC Neurodegeneration After Optic Nerve Crush in Mouse. Frontiers in cellular neuroscience 14, 109 (2020).

43. Li, L., et al. Single-cell transcriptome analysis of regenerating RGCs reveals potent glaucoma neural repair genes. Neuron 110, 2646–2663 e2646 (2022).

44. Chou, T.H., Bohorquez, J., Toft-Nielsen, J., Ozdamar, O. & Porciatti, V. Robust mouse pattern electroretinograms derived simultaneously from each eye using a common snout electrode. Invest Ophthalmol Vis Sci 55, 2469–2475 (2014).

45. Porciatti, V. Electrophysiological assessment of retinal ganglion cell function. Experimental eye research 141, 164–170 (2015).

46. Prusky, G.T., Alam, N.M., Beekman, S. & Douglas, R.M. Rapid quantification of adult and developing mouse spatial vision using a virtual optomotor system. Invest Ophthalmol Vis Sci 45, 4611–4616 (2004).

47. Douglas, R.M., et al. Independent visual threshold measurements in the two eyes of freely moving rats and mice using a virtual-reality optokinetic system. Vis Neurosci 22, 677–684 (2005).

48. Vong, L., et al. Leptin action on GABAergic neurons prevents obesity and reduces inhibitory tone to POMC neurons. Neuron 71, 142–154 (2011).

49. Tran, N.M., et al. Single-Cell Profiles of Retinal Ganglion Cells Differing in Resilience to Injury Reveal Neuroprotective Genes. Neuron 104, 1039–1055 e1012 (2019).

50. Jacobi, A., et al. Overlapping transcriptional programs promote survival and axonal regeneration of injured retinal ganglion cells. Neuron 110, 2625–2645 e2627 (2022).

51. Oswald, M.J., Tantirigama, M.L., Sonntag, I., Hughes, S.M. & Empson, R.M. Diversity of layer 5 projection neurons in the mouse motor cortex. Frontiers in cellular neuroscience 7, 174 (2013).

52. Scherrer, G., et al. VGLUT2 expression in primary afferent neurons is essential for normal acute pain and injury-induced heat hypersensitivity. Proc Natl Acad Sci U S A 107, 22296–22301 (2010).

53. Pivetta, C., Esposito, M.S., Sigrist, M. & Arber, S. Motor-circuit communication matrix from spinal cord to brainstem neurons revealed by developmental origin. Cell 156, 537–548 (2014).

54. Xu, Z., et al. Whole-brain connectivity atlas of glutamatergic and GABAergic neurons in the mouse dorsal and median raphe nuclei. eLife 10 (2021).

55. Xu, J., et al. Intersectional mapping of multi-transmitter neurons and other cell types in the brain. Cell reports 40, 111036 (2022).

56. Borgius, L., Restrepo, C.E., Leao, R.N., Saleh, N. & Kiehn, O. A transgenic mouse line for molecular genetic analysis of excitatory glutamatergic neurons. Molecular and cellular neurosciences 45, 245–257 (2010).

57. Lazarou, M., et al. The ubiquitin kinase PINK1 recruits autophagy receptors to induce mitophagy. Nature 524, 309–314 (2015).

58. Wong, Y.C. & Holzbaur, E.L. Optineurin is an autophagy receptor for damaged mitochondria in parkin-mediated mitophagy that is disrupted by an ALS-linked mutation. Proc Natl Acad Sci U S A 111, E4439–4448 (2014).

59. Evans, C.S. & Holzbaur, E.L. Degradation of engulfed mitochondria is rate-limiting in Optineurin-mediated mitophagy in neurons. eLife 9 (2020).

60. Ferree, A.W., et al. MitoTimer probe reveals the impact of autophagy, fusion, and motility on subcellular distribution of young and old mitochondrial protein and on relative mitochondrial protein age. Autophagy 9, 1887–1896 (2013).

61. Hernandez, G., et al. MitoTimer: a novel tool for monitoring mitochondrial turnover. Autophagy 9, 1852–1861 (2013).

62. Chertkova, A.O., et al. Robust and Bright Genetically Encoded Fluorescent Markers for Highlighting Structures and Compartments in Mammalian Cells. bioRxiv, 160374 (2020).

63. Lin, M.Y., et al. Releasing Syntaphilin Removes Stressed Mitochondria from Axons Independent of Mitophagy under Pathophysiological Conditions. Neuron 94, 595–610 e596 (2017).

64. Zhou, B., et al. Facilitation of axon regeneration by enhancing mitochondrial transport and rescuing energy deficits. J Cell Biol 214, 103–119 (2016).

65. Han, Q., et al. Restoring Cellular Energetics Promotes Axonal Regeneration and Functional Recovery after Spinal Cord Injury. Cell metabolism 31, 623–641 e628 (2020).

66. Luppi, P.H., Fort, P. & Jouvet, M. Iontophoretic application of unconjugated cholera toxin B subunit (CTb) combined with immunohistochemistry of neurochemical substances: a method for transmitter identification of retrogradely labeled neurons. Brain Res 534, 209–224 (1990).

67. Angelucci, A., Clasca, F. & Sur, M. Anterograde axonal tracing with the subunit B of cholera toxin: a highly sensitive immunohistochemical protocol for revealing fine axonal morphology in adult and neonatal brains. J Neurosci Methods 65, 101–112 (1996).

68. Branon, T.C., et al. Efficient proximity labeling in living cells and organisms with TurboID. Nature biotechnology 36, 880–887 (2018).

69. Xu, Y., Fan, X. & Hu, Y. In vivo interactome profiling by enzyme-catalyzed proximity labeling. Cell Biosci 11, 27 (2021).

70. Heo, J.M., et al. Integrated proteogenetic analysis reveals the landscape of a mitochondrial-autophagosome synapse during PARK2-dependent mitophagy. Sci Adv 5, eaay4624 (2019).

71. Henrichs, V., et al. Mitochondria-adaptor TRAK1 promotes kinesin-1 driven transport in crowded environments. Nature communications 11, 3123 (2020).

72. Han, S.M., Baig, H.S. & Hammarlund, M. Mitochondria Localize to Injured Axons to Support Regeneration. Neuron 92, 1308–1323 (2016).

73. Cartoni, R., et al. The Mammalian-Specific Protein Armcx1 Regulates Mitochondrial Transport during Axon Regeneration. Neuron 92, 1294–1307 (2016).

74. Huang, N., et al. Reprogramming an energetic AKT-PAK5 axis boosts axon energy supply and facilitates neuron survival and regeneration after injury and ischemia. Curr Biol 31, 3098–3114 e3097 (2021).

75. Kalinski, A.L., et al. Deacetylation of Miro1 by HDAC6 blocks mitochondrial transport and mediates axon growth inhibition. J Cell Biol 218, 1871–1890 (2019).

76. Chernyshova, K., Inoue, K., Yamashita, S.I., Fukuchi, T. & Kanki, T. Glaucoma-Associated Mutations in the Optineurin Gene Have Limited Impact on Parkin-Dependent Mitophagy. Invest Ophthalmol Vis Sci 60, 3625–3635 (2019).

77. Guillaud, L., El-Agamy, S.E., Otsuki, M. & Terenzio, M. Anterograde Axonal Transport in Neuronal Homeostasis and Disease. Frontiers in molecular neuroscience 13, 556175 (2020).

78. Shah, S.H. & Goldberg, J.L. The Role of Axon Transport in Neuroprotection and Regeneration. Developmental neurobiology 78, 998–1010 (2018).

79. Shah, S.H., et al. Quantitative transportomics identifies Kif5a as a major regulator of neurodegeneration. eLife 11 (2022).

80. Yokota, S., et al. Kif5a Regulates Mitochondrial Transport in Developing Retinal Ganglion Cells In Vitro. Invest Ophthalmol Vis Sci 64, 4 (2023).

81. Shlevkov, E., et al. A High-Content Screen Identifies TPP1 and Aurora B as Regulators of Axonal Mitochondrial Transport. Cell reports 28, 3224–3237 e3225 (2019).

82. Chi, Z.L., et al. Overexpression of optineurin E50K disrupts Rab8 interaction and leads to a progressive retinal degeneration in mice. Hum Mol Genet 19, 2606–2615 (2010).

83. Tseng, H.C., et al. Visual impairment in an optineurin mouse model of primary open-angle glaucoma. Neurobiology of aging 36, 2201–2212 (2015).

84. Ito, Y., et al. RIPK1 mediates axonal degeneration by promoting inflammation and necroptosis in ALS. Science 353, 603–608 (2016).

85. Buffelli, M., et al. Genetic evidence that relative synaptic efficacy biases the outcome of synaptic competition. Nature 424, 430–434 (2003).

86. Fang, F., et al. NMNAT2 is downregulated in glaucomatous RGCs, and RGC-specific gene therapy rescues neurodegeneration and visual function. Mol Ther 30, 1421–1431 (2022).

87. Masin, L., et al. A novel retinal ganglion cell quantification tool based on deep learning. Scientific reports 11, 702 (2021).

88. Renier, N., et al. iDISCO: a simple, rapid method to immunolabel large tissue samples for volume imaging. Cell 159, 896–910 (2014).

89. Vowinckel, J., Hartl, J., Butler, R. & Ralser, M. MitoLoc: A method for the simultaneous quantification of mitochondrial network morphology and membrane potential in single cells. Mitochondrion 24, 77–86 (2015).

90. Basu, H., Ding, L., Pekkurnaz, G., Cronin, M. & Schwarz, T.L. Kymolyzer, a Semi-Autonomous Kymography Tool to Analyze Intracellular Motility. Curr Protoc Cell Biol 87, e107 (2020).

91. Wang, X. & Schwarz, T.L. The mechanism of Ca2+ -dependent regulation of kinesin-mediated mitochondrial motility. Cell 136, 163–174 (2009).

92. Chen, W.W., Freinkman, E. & Sabatini, D.M. Rapid immunopurification of mitochondria for metabolite profiling and absolute quantification of matrix metabolites. Nature protocols 12, 2215–2231 (2017).

93. Bayraktar, E.C., et al. MITO-Tag Mice enable rapid isolation and multimodal profiling of mitochondria from specific cell types in vivo. Proc Natl Acad Sci U S A 116, 303–312 (2019).

94. 94. Laser peripheral iridotomy for pupillary-block glaucoma. American Academy of Ophthalmology. Ophthalmology 101, 1749-1758 (1994).

95. 95. Schindelin, J., et al. Fiji: an open-source platform for biological-image analysis. Nature methods 9, 676-682 (2012).

96. Jumper, J., et al. Highly accurate protein structure prediction with AlphaFold. Nature 596, 583–589 (2021).

97. 97. Evans, R., et al. Protein complex prediction with AlphaFold-Multimer. bioRxiv, 2021.2010.2004.463034 (2022).

