## Supplementary Data for "Optineurin-facilitated axonal mitochondria delivery promotes neuroprotection and axon regeneration"

**The PDF file includes:**

Figs. S1 to S7

**Other Supplementary Materials for this manuscript include the following:**

Movies S1 to S7

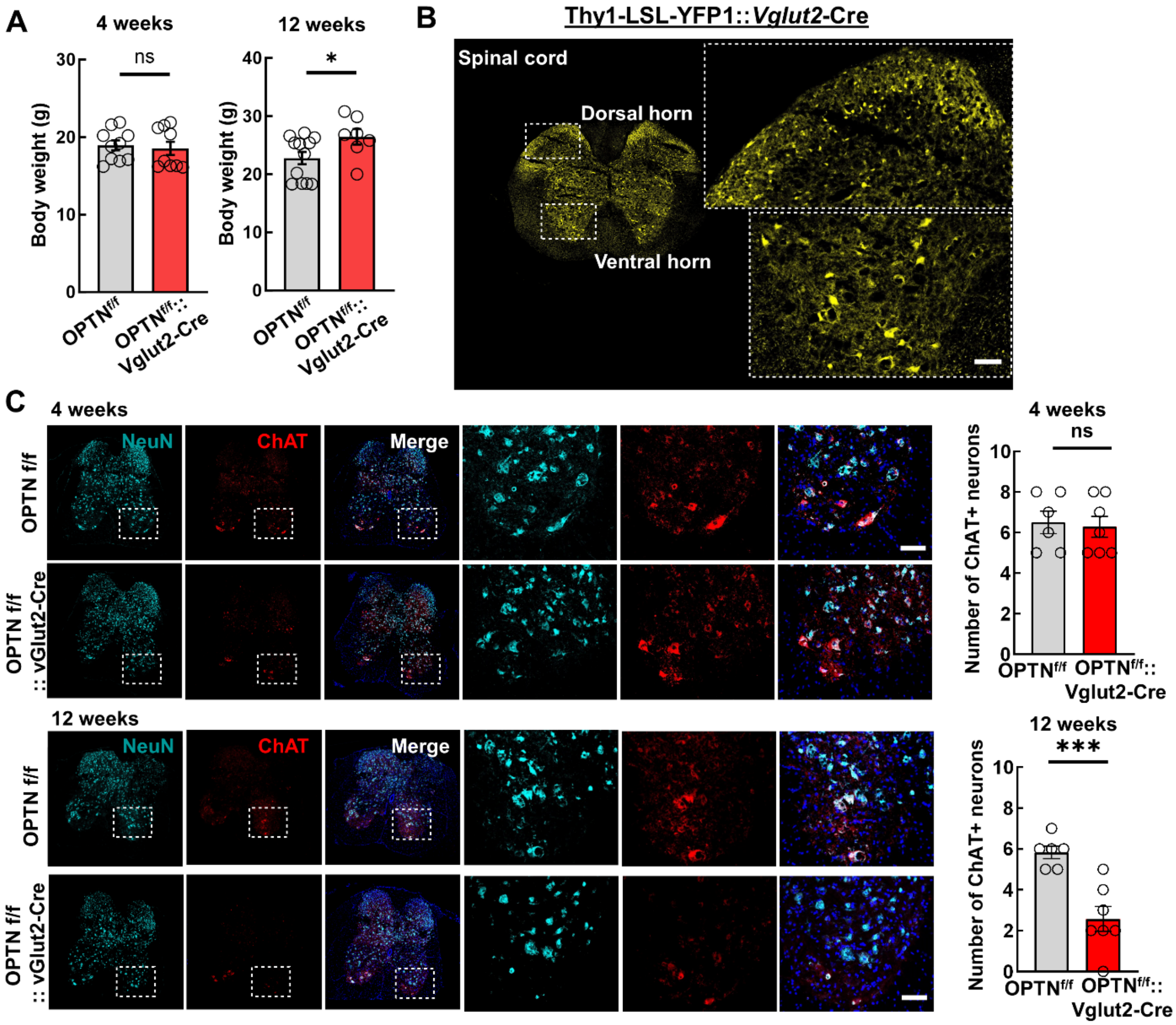


**Fig. S1. Vglut2-Cre mediated OPTNΔC in neurons causes body weight change and motor neuron degeneration in OPTN$^{f/f}$::Vglut2-Cre mice, related to Figure 2. A,** Body weight measurements at 4w and 12w. n = 9-12 mice. **B,** Representative confocal images of spinal cord sections in the Thy1-LSL-YFP-1::Vglut2-Cre mice. Higher magnification images of spinal cord dorsal and ventral horns are shown to the right. Scale bar, 50 μm. **C,** Representative confocal images of lumbar spinal cord sections coimmunostained with NeuN and ChAT for motor neuron and DAPI for cell nuclei of 4 weeks and 12 weeks old OPTNf/f::Vglut2-Cre mice. Enlarged images of framed regions in the ventral horns are shown to the right. Scale bar, 50 μm. Quantification of

ChAT-positive motor neuron survival at 4 or 12 weeks old are shown in the right panel. $n = 6$-7 mice. All the quantification data are presented as means ± s.e.m, *: $p<0.05$, ***: $p < 0.001$, ns: no significance, unpaired Student's t-test.

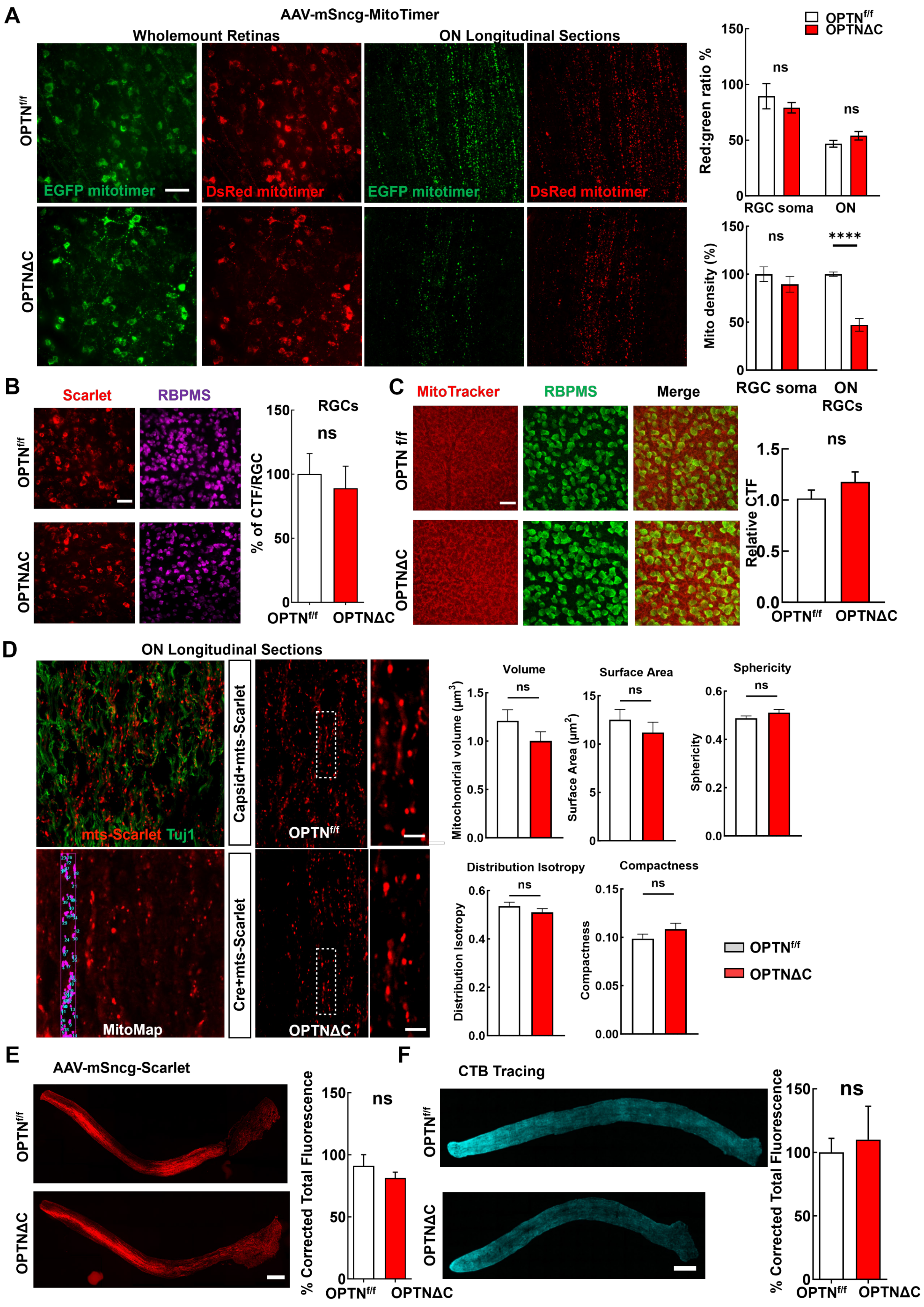
A
AAV-mSncg-MitoTimer
Wholemount Retinas
ON Longitudinal Sections
OPTNf/f
OPTNΔC
EGFP mitotimer
DsRed mitotimer
Red:green ratio %
RGC soma
ON
Mito density (%)
ns
****
B
Scarlet
RBPMS
RGCs
% of CTF/RGC
C
MitoTracker
RBPMS
Merge
OPTN f/f
Relative CTF
D
ON Longitudinal Sections
mts-Scarlet Tuj1
Capsid+mts-Scarlet
Cre+mts-Scarlet
MitoMap
Volume
Surface Area
Sphericity
Mitochondrial volume (μm3)
Surface Area (μm2)
Distribution Isotropy
Compactness
E
AAV-mSncg-Scarlet
% Corrected Total Fluorescence
F
CTB Tracing

**Fig. S2. OPTNΔC significantly decreases ON mitochondrial density but does not significantly affect mitophagy, RGC mitochondrial density, mitochondrial morphology, or general axonal transportation, related to Figure 3. A,** Representative images of retina wholemounts and ON longitudinal sections 2 weeks after intravitreal injection of AAV-Cre + AAV-MitoTimer in $OPTN^{f/f}$ mice, Scale bar, 50 µm. Quantification of red to green fluorescence intensity ratio and mitochondrial density of RGC somata in retinas and axons in ONs. *n* = 5 mice. **B,** Representative images of retina wholemount 2 weeks after intravitreal injection of AAV-Cre + AAV-4xMTS-Scarlet in $OPTN^{f/f}$ mice. Scale bar, 50 µm. Quantification of Corrected Total Fluorescence (CTF)/RGC, represented as a percentage of OPTNΔC eyes compared to the contralateral naïve $OPTN^{f/f}$ (CL) eyes. *n* = 5 mice. **C,** Representative images of MitoTracker labeled-retinal wholemounts. Scale bar, 50 µm. Quantification of CTF, represented as a ratio of OPTNΔC eyes compared to the CL eyes. *n* = 5 mice. **D,** Representative images of ON longitudinal sections 2 weeks after intravitreal injection of AAV-Cre + AAV-4xMTS-Scarlet in $OPTN^{f/f}$ mice. Axons are immunostained with Tuj1 antibody. Higher magnification images are shown to the right. Scale bar, 10 µm. MitoMap analysis of mitochondrial volume, surface area, sphericity, distribution isotropy and compactness does not show significant difference between $OPTN^{f/f}$ and OPTNΔC ONs. *n* = 3 mice. **E,** Representative images of ON longitudinal sections 2 weeks after intravitreal injection of AAV-Cre + AAV-Scarlet. Scale bar, 200 µm. Quantification of total Scarlet fluorescence, represented as a percentage of OPTNΔC eyes compared to the CL eyes. *n* = 4 mice. **F,** Representative images of ON longitudinal sections 2 weeks after intravitreal injection of AAV-Cre and 3 days post-injection with Cholera Toxin subunit B (CTB) Alexa Fluor-555 conjugate. Scale bar, 200 µm. Quantification of total CTB fluorescence, represented as a percentage of

OPTNΔC eyes compared to the CL eyes. *n* = 3 mice. All the quantification data are presented as means ± s.e.m, ns, no significance, ∗∗∗∗: $p < 0.0001$, Student's t-test.

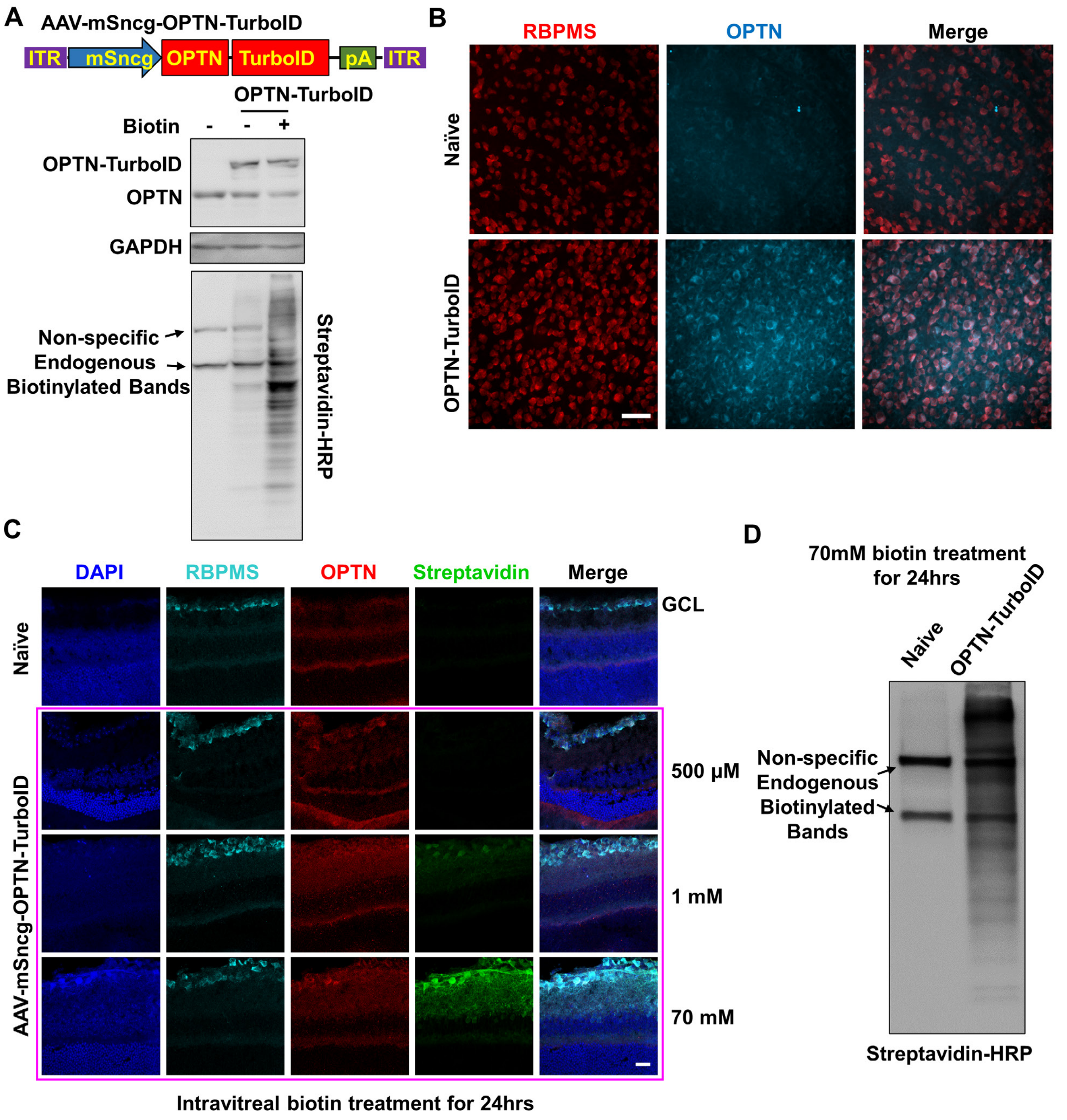


**Fig. S3.** ***In vivo*** **RGC TurboID assay development, related to Figure 3E. A**, (top) AAV vector to drive human OPTN-TurboID expression in RGCs. (bottom) Naïve and OPTN-TurboID expressing HEK293 cells were treated with or without 500µM biotin for 24hrs. OPTN-TurboID expression and OPTN-TurboID-mediated protein biotinylation in HEK293 cells identified by

Western blotting. **B**, Representative images of retina wholemounts demonstrating AAV-mediated OPTN-TurboID overexpression in RGCs. Scale bar, 50 µm. **C**, Representative images of retina sections with OPTN-TurboID expression 24 hours after intravitreal delivery of various amounts of biotin. Scale bar, 20 µm. **D**, Western blotting of retina lysates 24 hours after intravitreal delivery of 70mM biotin, demonstrating biotinylated proteins in RGCs with HRP-conjugated streptavidin.

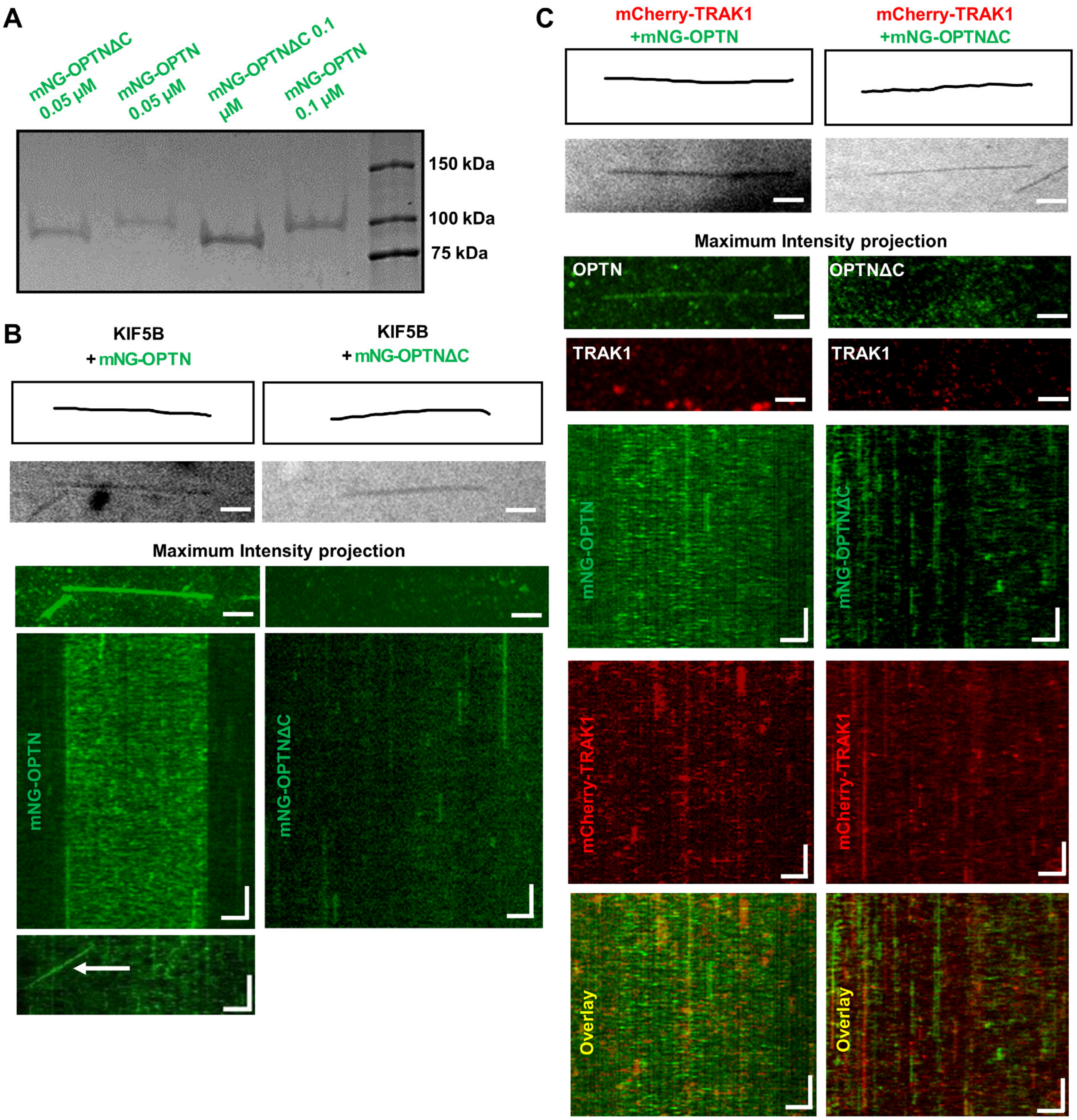


**Fig. S4. OPTN binds to microtubules in a C-terminus dependent manner, which is not affected by TRAK1 or KIF5B alone, related to Figure 4. A,** SDS gel showing purified mNG-OPTN and mNG-OPTNΔC. **B**, (top to bottom) IRM image of microtubules, maximum intensity projection of mNG-OPTN/mNG-OPTNΔC, Kymographs of 0.1 µM mNG-OPTN or 0.1 µM mNG-OPTNΔC binding and unbinding to microtubules in the presence of 1 nM unlabeled KIF5B (representative rare migration event marked by arrow). Horizontal scale bar = 2 μm, Vertical scale bar = 4 seconds. *n* = 3 experiments. **C**, (top to bottom) RM image of microtubules, maximum intensity projection of mNG-OPTN/mNG-OPTNΔC and TRAK1-mCherry, Kymograph of 10 nM OPTN / 30 nM OPTNΔC binding and unbinding to microtubules in the presence of 17 nM TRAK1-mCherry. Horizontal scale bar = 2 μm, Vertical scale bar = 8 seconds. *n* = 3 experiments.

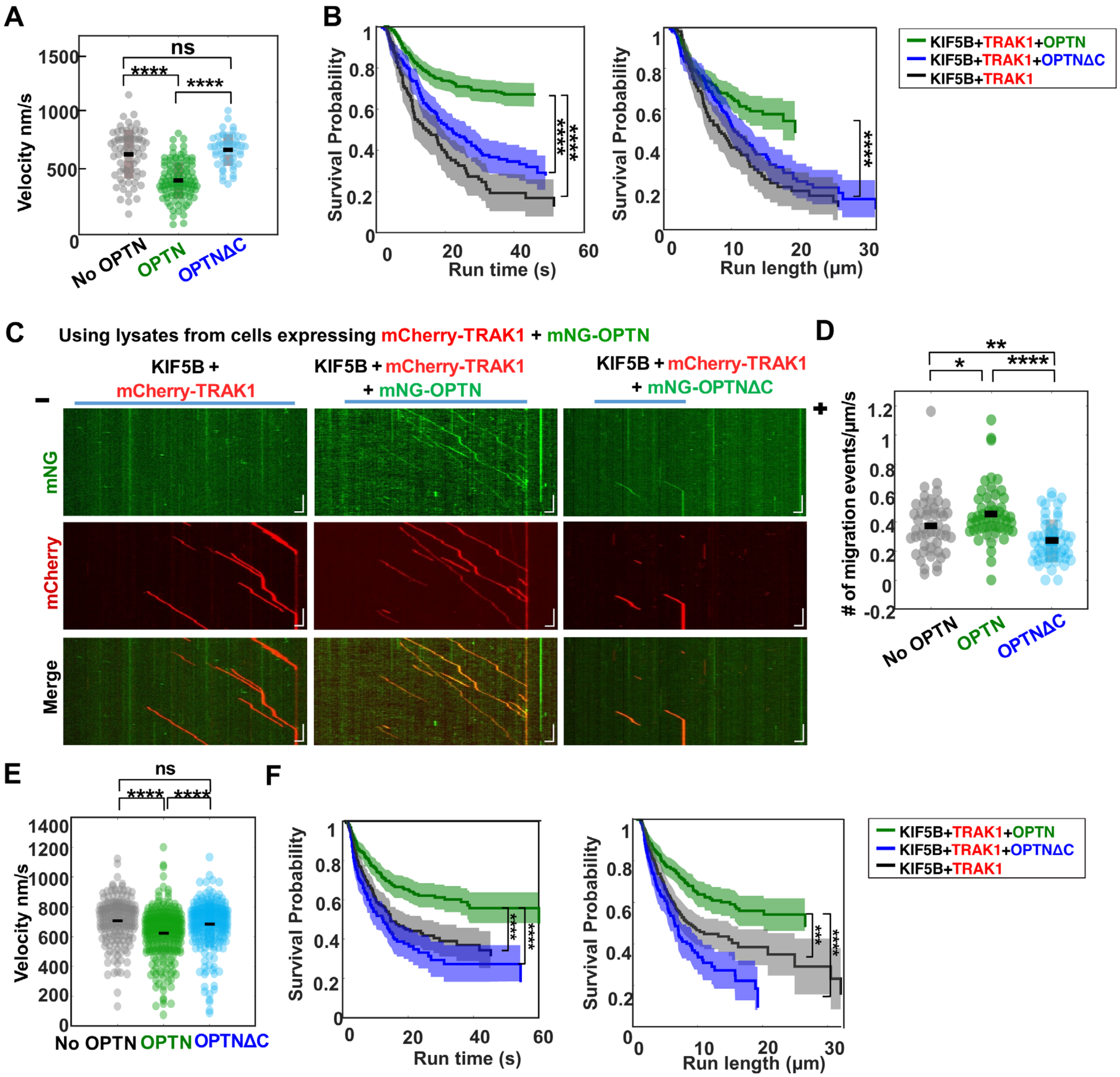


**Fig. S5. *In vitro* motility assay of immobilized microtubules with recombinant proteins or cell lysates expressing mNG-OPTN and TRAK1-mCherry and *ex vivo* ON mitochondria kymograph assays, related to Figure 4. A,** Velocity of continuous migration events of complexes of KIF5B-TRAK1-OPTN (n = 101), KIF5B-TRAK1-OPTNΔC (n = 50), and KIF5B-TRAK1 (n = 68), respectively. *n* = 3 experiments, ****: p $<10^{-13}$, ns = 0.267, t-test. **B,** Left, Run time probability distribution of complexes of KIF5B-TRAK1-OPTN (green, n = 332), KIF5B-TRAK1-

OPTNΔC (blue, n = 208), and KIF5B-TRAK1 (black, n = 118) on microtubules. Shaded regions indicate the 95% confidence intervals. Right, Run length probability distribution (Kaplan-Meir estimation) of complexes of KIF5B-TRAK1-OPTN (green, n = 332), KIF5B-TRAK1-OPTNΔC (blue, n = 208) and KIF5B-TRAK1 (black, n = 118) on microtubules. Shaded regions indicate the 95% confidence intervals. *n* = 4-6 experiments, ****: $p < 0.0001$, t-test. **C,** *In vitro* motility assay of immobilized microtubules with cell lysate expressing TRAK1-mCherry and mNG-OPTN. Kymograph of TRAK1 with OPTN or OPTNΔC in the presence of unlabeled KIF5B walking to the plus end of microtubules. Horizontal scale bar, 2 μm; vertical scale bar, 10 seconds. Blue bars indicate the microtubule positions along the kymograph. **D,** Frequency of migration events (/μm/s) of complexes of KIF5B-TRAK1 (n =48), KIF5B-TRAK1-OPTN (n = 52), KIF5B-TRAK1-OPTNΔC (n = 52), N = 4 experiments. unpaired Student's t-test. **E,** Velocity of continuous migration events of complexes of KIF5B-TRAK1 (n =236), KIF5B-TRAK1-OPTN (n =274) and KIF5B-TRAK1-OPTNΔC (n =177), respectively. N = 4 experiments, unpaired Student's t-test. **F,** Left, Run time probability distribution of complexes of KIF5B-TRAK1-OPTN (green, n = 327, N=4), KIF5B-TRAK1-OPTNΔC (blue, n = 248, N=4), and KIF5B-TRAK1 (black, n = 220, N=4) on microtubules. Shaded regions indicate the 95% confidence intervals. Right, Run length probability distribution (Kaplan-Meir estimation) of complexes of KIF5B-TRAK1-OPTN (green, n = 327, N=4), KIF5B-TRAK1-OPTNΔC (blue, n = 248, N=4) and KIF5B-TRAK1 (black, n = 220, N=4) on microtubules. Shaded regions indicate the 95% confidence intervals. n=number of molecules, N= number of independent trails. **D-F**, ∗: $p < 0.05$, ∗∗: $p < 0.01$, ∗∗∗: $p < 0.001$, ∗∗∗∗: $p < 0.0001$, ns: no significance, with t-test.

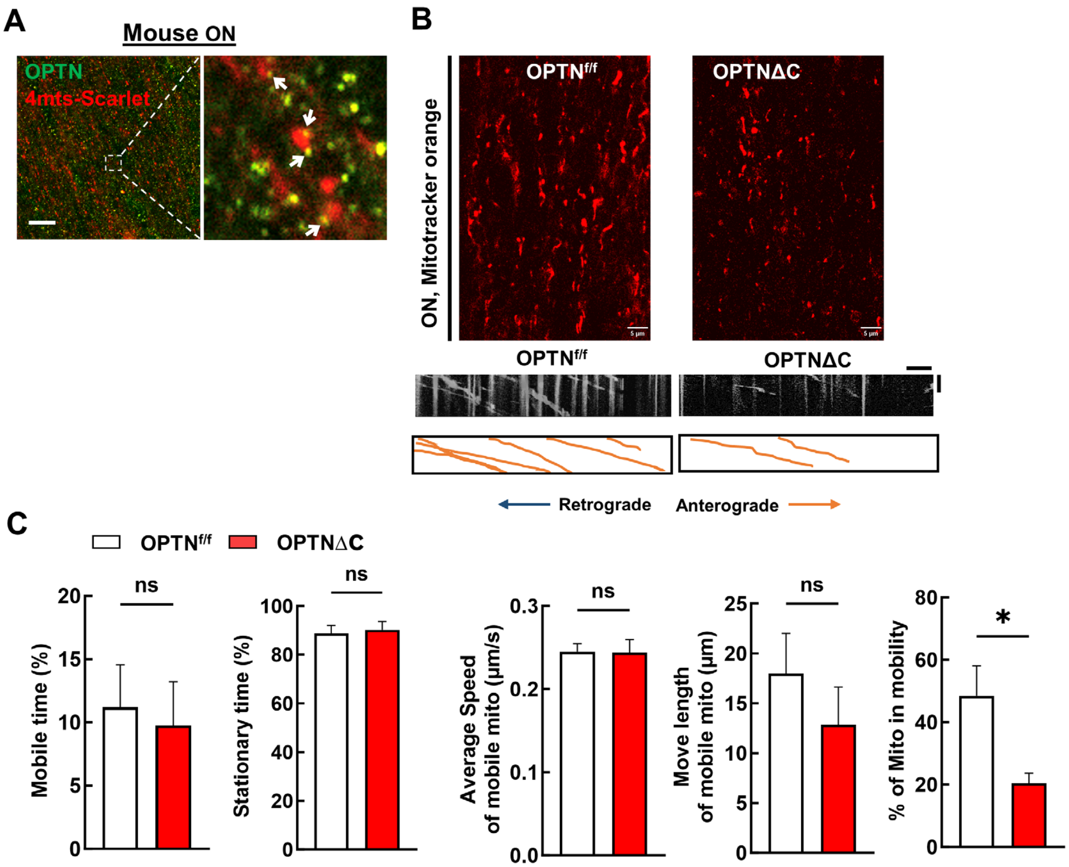


**Fig. S6. *Ex vivo* time-lapse imaging revealed mitochondrial trafficking deficits in OPTNΔC-optic nerves. A,** Immunostaining of endogenous OPTN in mouse ONs with mitochondria labeled with 4MTS-Scarlet. White arrows indicate the OPTN punctas on the surfaces of mitochondria. Scale bar, 20 µm. **B,** Upper, Representative ON wholemount images of the OPTN$^{f/f}$ and OPTNΔC mice with MitoTracker Orange labeling, Scale bars, 20 µm. Lower, Kymograph and traces of MitoTracker labeled mitochondria movement along the axons in the ONs of the OPTN$^{f/f}$ and OPTNΔC mice. Horizontal scale bar, 5 µm; vertical scale bar, 1 minute. **C,** Quantification of each mitochondrion time in motion and time in stationary, average speed and move length of each mobile mitochondrion, and percentage of mitochondria in motion. n = 21–30 mitochondria from

3 axons per group. Data are presented as means ± s.e.m, *: $p<0.05$, ns, no significance, with Student's t-test.

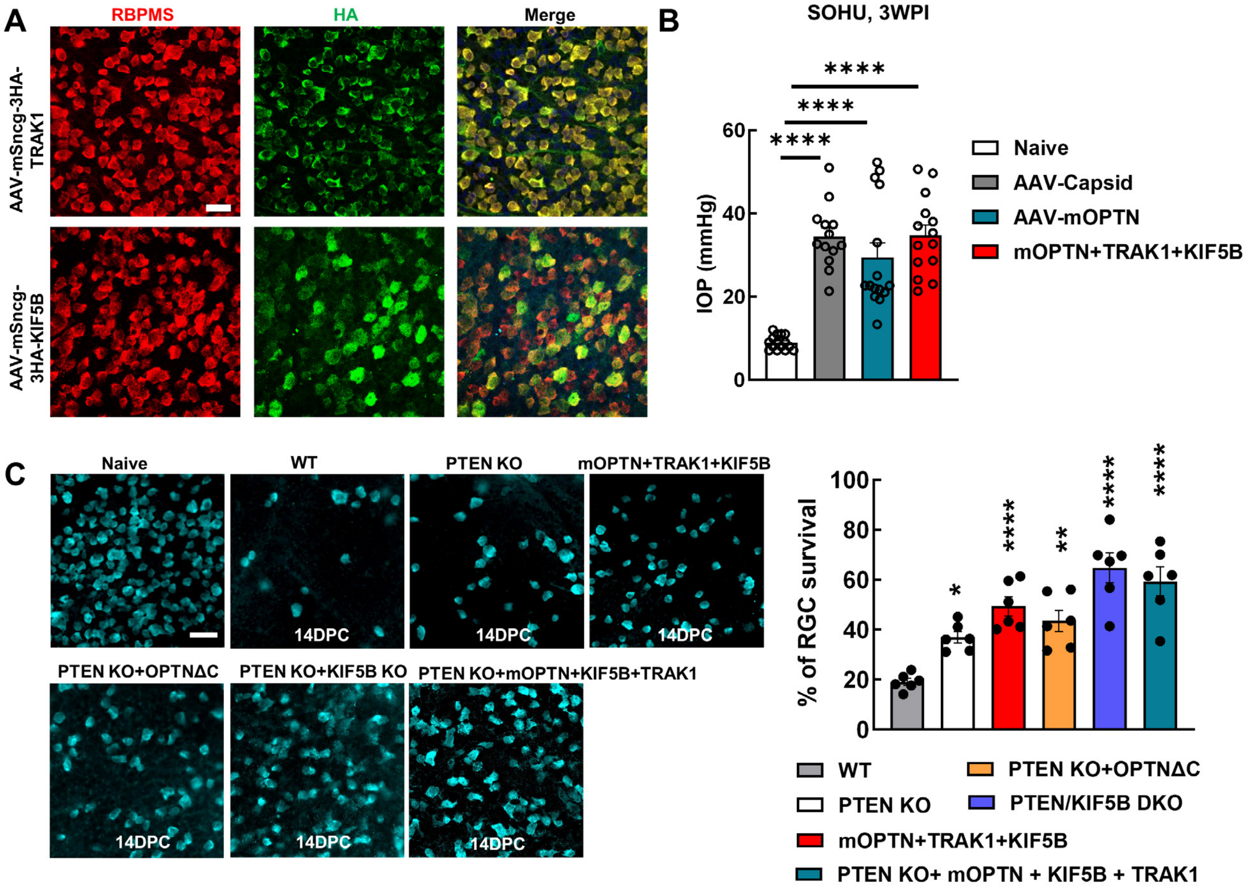


**Fig. S7. AAV-mediated KIF5B and/or TRAK1 overexpression in RGCs promote RGC survival, related to Figures 5-7. A**, Representative confocal images of retinal wholemounts showing AAV2-mSncg promoter-mediated TRAK1 and KIF5B expression in RBPMS-positive RGCs 2 weeks post AAV intravitreal injection. Scale bar, 20 µm. **B,** IOP of Naïve and SOHU eyes from different groups of mice at 3wpi. n = 13-15 mice. Data are presented as means ± s.e.m, ****: p<0.0001, with one-way ANOVA and *post hoc* Dunnett's comparison test. **C,** Left, representative confocal images of the retinal wholemounts showing surviving RBPMS-positive RGCs at 14dpc, Scale bar, 50 µm. Right, quantification of surviving RGC somata in peripheral retina at 14dpc, represented as percentage of crushed eyes compared to the CL eyes. Data are presented as means ± s.e.m, n = 6 in each group. *: p<0.05, **: p<0.01, ****: p<0.0001, one-way ANOVA with *post hoc* Dunnett's comparison test.

**Movie S1. OPTN binds to microtubules in a C-terminus dependent manner.** Time-lapse imaging of purified mNG-OPTN or mNG-OPTNΔC binding and unbinding to immobilized microtubules. Scale bar, 2.1 μm.

**Movie S2. Lysates of OPTN expressing cells bind to microtubules in a C-terminus dependent manner.** Time-lapse imaging of lysates from cells expressing mNG-OPTN or mNG-OPTNΔC binding and unbinding to immobilized microtubules. Scale bar, 2 μm.

**Movie S3. TRAK1-KIF5B migration on microtubules with or without OPTN or OPTNΔC from purified proteins.** Time-lapse imaging of purified mCherry-TRAK1 walking to the plus ends of immobilized microtubules in the presence of unlabeled KIF5B with or without purified mNG-OPTN or mNG-OPTNΔC. Scale bar, 2 μm.

**Movie S4. TRAK1-KIF5B migration on microtubules with or without OPTN or OPTNΔC from cell lysates.** Time-lapse imaging of lysates from cells expressing mCherry-TRAK1 with or without mNG-OPTN or mNG-OPTNΔC walking to the plus ends of immobilized microtubules in the presence of unlabeled KIF5B. Scale bar, 2 μm.

**Movie S5. Mitochondria migration in cultured hippocampal neuron axons in the presence of OPTN or OPTNΔC.** Time-lapse imaging of mitochondria (labeled with MitoDsRed) showing anterograde movement to the right and retrograde movement to the left in $OPTN^{f/f}$ or OPTNΔC hippocampal neurons. Axons are labelled with AAV-mSncg-EGFP ($OPTN^{f/f}$) or AAV-mSncg-Cre-T2A-EGFP (OPTNΔC). Scale bar, 10 μm,

**Movie S6. Mitochondria migration in *ex vivo* ONs in the presence of OPTN or OPTNΔC.** Time-lapse imaging of mitochondria (labeled with MitoTracker Orange) movements along the axons in *ex vivo* ONs of the $OPTN^{f/f}$ and OPTNΔC mice. Scale bar, 5 µm.

**Movie S7. Mitochondria migration in *ex vivo* ONs of naïve, glaucomatous, and treated glaucomatous mice.** Time-lapse imaging of mitochondria (labeled with MitoTracker Orange) movements along the axons in *ex vivo* ONs of the naïve, SOHU glaucoma 1wpi, and SOHU glaucoma mice treated with mOPTN+TRAK1+KIF5B. Scale bar, 20 µm.
